# Centrosome-dependent microtubule organization sets the conditions for axon formation

**DOI:** 10.1101/2020.12.29.424696

**Authors:** Durga Praveen Meka, Oliver Kobler, Souhaila Wuesthoff, Birgit Schwanke, Christoph Krisp, Nessa Schmuelling, René Rueter, Tabitha Ruecker, Hartmut Schlüter, Eugenio F. Fornasiero, Froylan Calderon de Anda

## Abstract

Microtubule remodeling is critical during axon development when the more stable microtubules populate the axon. It is not completely understood, however, how this local cytoskeleton remodeling is coordinated. The centrosome, the main microtubule-organizing center (MTOC), has been suggested to be crucial for axon specification ^1–5^. Conversely, it was proposed that axon elongation is independent of centrosomal functions ^6^. Here we report that microtubule dynamics in early neurons follow a radial organization which establishes the conditions for the axon formation. Using high-resolution microscopy of early developing neurons, we demonstrate that few somatic acetylated microtubules are restricted near the centrosome. At later stages, however, acetylated microtubules spread out in the soma and concentrate in the growing axon. Furthermore, live-imaging of the microtubule plus-end binding protein EB3 in early differentiating neurons shows that growing microtubules have increased length and growth speed near the MTOC, suggesting local differences that might favor axon selection. Importantly, due to the lack of somatic stable/acetylated microtubules in early developing neurons, disruption of the F-actin cytoskeleton does not induce multiple axons, as it does at later stages of differentiation. Finally, we demonstrate that overexpression of the centrosomal protein 120 (Cep120), known for promoting microtubule acetylation and stabilization, induces multiple axons, while its downregulation decreases the content of proteins regulating microtubule dynamics and stability, hence hampering axon formation. Collectively, our data show that early centrosome-dependent microtubule organization contributes to axon formation.

## Introduction

Neurons are complex cells with distinct functional domains for information processing. In the postsynaptic compartment dendrites receive the synaptic input and relay signals to the soma, where the integrated information is transmitted via the axon over short or long distances. Differentiating neurons follow an intricate process during which one of the neurites is specified as an axon and the remaining typically become dendrites. Axon specification is the defining step for neuronal polarization. On this line, it has been shown that local actin instability in the growth cones or microtubule stability in the neurite shafts sustain axon elongation ^7, 8^. The neurite with more dynamic F-actin in the growth cone and more microtubules that are eventually more stable, develops as an axon, whereas the remaining neurites become dendrites ^7–12^. In cultured rat hippocampal neurons, the future axon was suggested to have more stable microtubules in its shaft, due to the enrichment of acetylated microtubules ^8^. Accordingly, global application of the microtubule-stabilizing drug Taxol, induced the formation of multiple axons. Moreover, microtubule stabilization increased F-actin dynamics in growth cones ^12^. However, the question of how microtubule acetylation enrichment occurs preferentially in the growing axon remains elusive.

To understand the role of microtubule stabilization during axon selection, it is crucial to consider if this process is intrinsically mediated or sustained through external cues. Several *in vitro* and *in situ* studies suggest key roles of centrosome-dependent radial microtubule organization in migrating and early differentiating neurons ^5, 13–17^. Consequently, inhibition of microtubule nucleation at the centrosome hinder microtubule reassembly and compromise axonal growth ^18^, attributing an important role for the centrosome and its microtubule assembling ability in axon formation. In support of this idea, the first neurite in a newly formed neuron correlates with the position of the centrosome and the Golgi apparatus. Eventually the first emerged neurite becomes the axon ^1, 2^. It is not clear, however, whether the centrosome plays a role for the stabilization/acetylation of microtubules in the growing axon. To test this possibility, we performed a series of state-of-the-art experiments including super-resolution imaging, differential quantitative proteomics and high-content time-lapse microscopy to examine microtubule organization in early developing neurons.

Mechanistically, we show that Cep120, a centriolar protein which induces microtubule acetylation and stabilization during cilia formation ^19–21^, modulates microtubules dynamics and axon formation. Thus, Cep120 overexpression produces neurons with multiple axons. Overall, our data show a radial organization of microtubules which favors preferential microtubules stability and thus axon formation.

## Results

### Radial organization of acetylated microtubules in early developing neurons

Axon formation is a hallmark of neuronal polarization in early developing hippocampal and cortical pyramidal neurons ^4, 22–25^. Neurons initially extend several neurites (Stage 2; ^22^), from which usually those with the fastest growth rate become the axon (Stage 3; ^22^), while the remaining neurites transform into dendrites ^22, 26^. It is known that during the transition from stage 2 to 3 the growth cone of the neurite, which elongates as an axon, contains more dynamic F-actin ^7^ and more stable microtubules ^8^. In accord to previous data of others and our own, we found that in neurons without neurites (stage 1) the position of the centrosome is a landmark to predict the position of axon outgrowth ^1–4^. It is not clear, however, if the centrosome contributes with stable microtubules in the growing axon at stage 3. To explore this possibility, we decided to perform a detailed analysis the distribution of acetylated tubulin and to track stable microtubules in neurons from stage 1 to 3. To this end, we used confocal and super-resolution microscopy during early neuronal differentiation *in vitro.* Tyrosinated and acetylated tubulin were visualized in fixed cells via confocal and STimulated Emission Depletion (STED) microscopy. STED microscopy images revealed that stage 1 neurons have the acetylated tubulin concentrated around the centrosome (labeled with pericentrin) while the tyrosinated tubulin (unstable microtubules) spread out in the cell body (Figure 1a, b and Video 1). At stage 2, however, the acetylated tubulin localizes throughout the cell body, as the tyrosinated tubulin does, and start to penetrate the neurites (Figure 1a, b). Neurons which started to extend an axon (stage 3) increased the content of acetylated tubulin drastically in the longest neurite presumably the future axon (Figure 1c, d). These results suggest that the microtubule stabilization spreads radially from the centrosomal area to the growing neurites, eventually concentrating specifically in the longest neurite or future axon.

**Figure 1.**
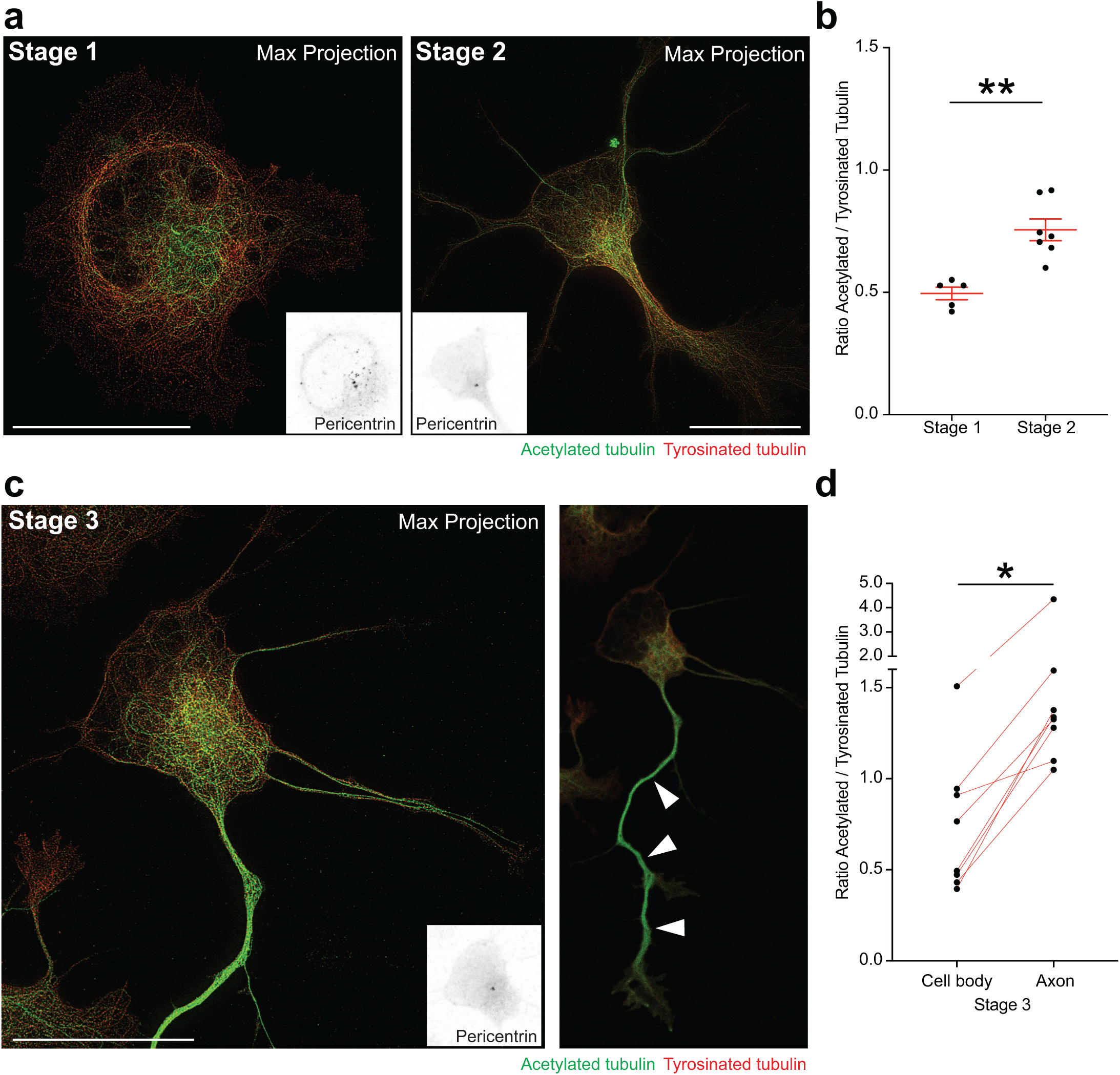
Acetylation of microtubules spreads radially from the centrosomal area towards the growing neurites before polarization and significantly in to the growing axon after symmetry breakage. **a.** STED images of acetylated and tyrosinated tubulin immunostaining in stage 1 and stage 2 hippocampal neurons. Insets: Centrosome labelled by Pericentrin antibody staining. Scale bar: 10 µm **b.** Quantifications show ratio of acetylated to tyrosinated tubulin signal in the soma of stage 1 and stage 2 neuronal soma. Mean ± SEM values for stage 1 neurons = 0.4954 ± 0.0255 and stage 2 neurons = 0.7556 ± 0.0449. ** P = 0.0011 by unpaired Student’s t-test. n = 5 - 7 cells from 2 different cultures. **c.** STED images of acetylated and tyrosinated tubulin immunostaining in a stage 3 hippocampal neuron Inset: centrosome labelled by Pericentrin antibody staining. White arrowheads point enrichment of tubulin acetylation in the axon. Scale bar: 10 µm **d.** Quantifications show ratio of acetylated to tyrosinated tubulin signal in stage 3 neuronal soma and stage 3 axons. Mean ± SEM values for stage 3 soma = 0.7400 ± 0.1341 and stage 3 axons =1.676 ± 0.3861. * P = 0.0133 by paired Student’s t-test. n = 8 cells from 2 different cultures.

### Microtube dynamics at the soma differ through the early neuronal development stages

To gain further insight into the relevance of this early somatic microtubule organization, we decided to monitor growing microtubules at the soma of developing neurons. Neurons were transfected with the microtubule plus-end marker tdTomato-EB3 before plating and 18-36hr later developing neurons (stage 1 to early stage 3) were imaged for 5 minutes with a frame rate of 2 seconds. To quantify microtubule dynamics, we performed semi-automatic tracking of EB3 particles (see Material and Methods for detailed description) to reconstitute microtubule tracks and quantified for each microtubule track the growth speed, displacement of EB3 comet per each frame, and growth lifetimes (Figure 2a; Videos 2, 3 and 4). We plotted the histograms of these parameters and calculated the median of the distribution for the EB3 tracks speed and fitted the EB3 comet displacement, lifetime with exponential decays (^27, 28^; Figure 2b). Additionally, we also plotted all events registered per cell (EB3 tracks speed, EB3 comet displacement, and growth lifetime; Supp Figure 1-3a, b). Our results show that the median growth speed of microtubules is increased at stage 2 cells compared to stages 1 and 3 (Figure 2b and Supp Figure 1a, b). The characteristic growth (λ) per frame is higher at stage 2 neurons compared to stages 1 and 3 (Figure 2b and Supp Figure 2a, b). However, the characteristic growth lifetime (τ) is higher at stage 3 compared to stages 1 and 2 (Figure 2b and Supp Figure 3a, b). These results highlight the highly dynamic nature of the somatic microtubules during neuronal polarization. Specifically, somatic microtubules underwent drastic changes before axon extension, at stage 2.

**Figure 2.**
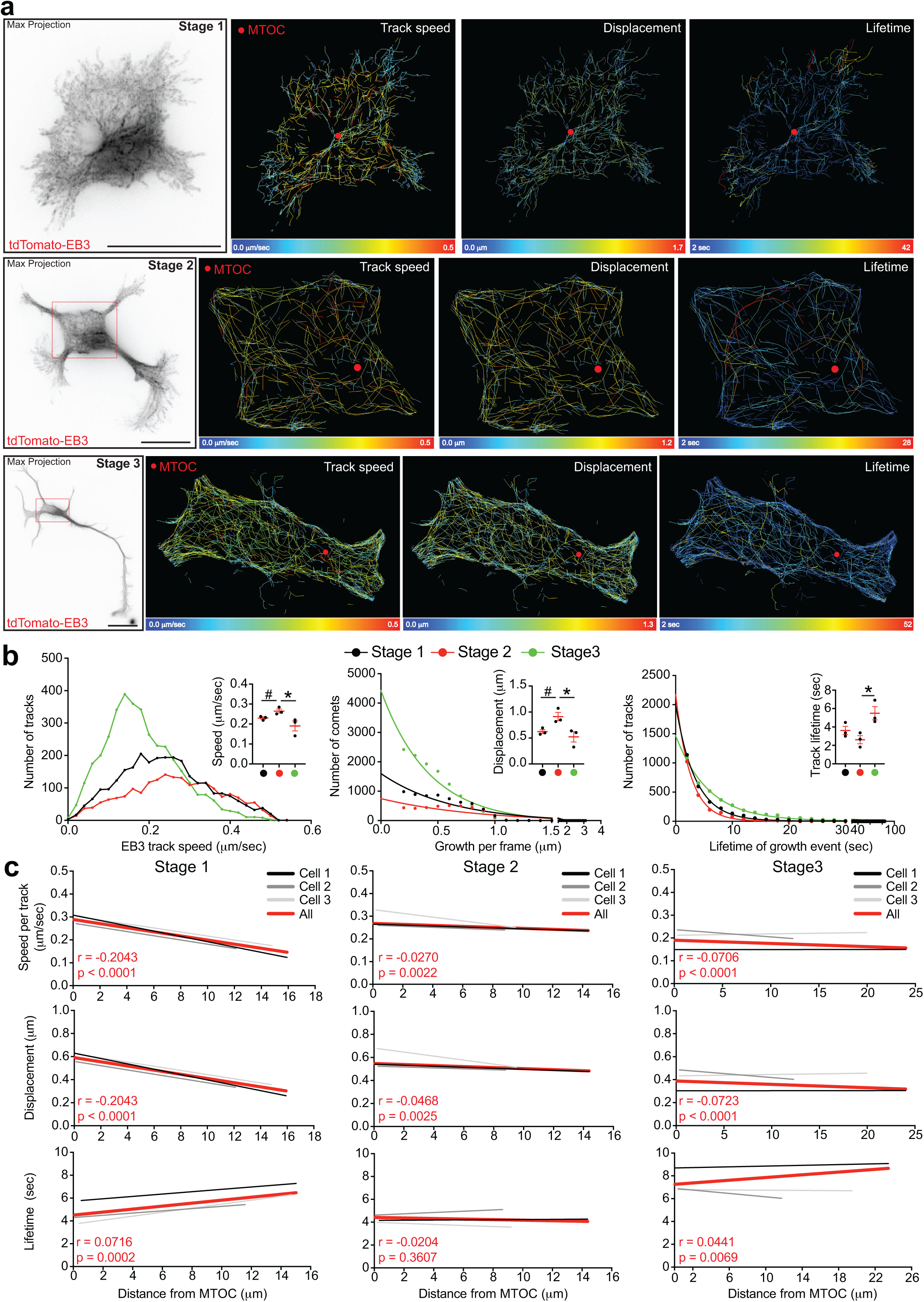
Microtubule plus-end (EB3) dynamics in the soma of developing neurons decrease during polarization: order of magnitude of EB3 dynamics = stage 1 (near MTOC in the soma) > stage 2 cells soma > stage 3 cells soma. **a.** Maximum intensity projection images (black on white images) from a 5 min time-lapse (1 frame every 2 seconds) of EB3-tdTomato-trasfected stage 1, stage 2 and stage 3 hippocampal neurons. EB3 track growth speed, EB3 displacement (per frame) and lifetime of the EB3 track analyzed by TrackMate ImageJ plugin from the respective neurons shown as heatmaps. Red dots in the heatmaps indicate MTOC. Scale bar: 10 µm. **b.** Left panel: Frequency distribution compares speed (µm/sec) of EB3 tracks in soma of stage 1, stage 2 and stage 3 neurons. Inset shows values of median speed (µm/sec). Mean ± SEM values of median speed from stage 1 = 0.2300 ± 0.0056, stage 2 = 0.2650 ± 0.0102, stage 3 = 0.1903 ± 0.02491. P = 0.0436 by one-way ANOVA, post hoc Tukey’s test, * P = 0.0367. For Stage 1 vs Stage 2: # P = 0.0406 by unpaired Student’s t-test. n = 3 cells per stage. Middle panel: Nonlinear fit histogram compares displacement of EB3 (in µm) per each frame (2sec) soma of stage 1, stage 2 and stage 3 cells. Inset shows characteristic growth (λ) per frame, which is the half values of displacement per frame. Mean ± SEM values of the characteristic growth (λ) per each frame for stage 1 = 0.6216 ± 0.03705, stage 2 = 0.9120 ± 0.08048, stage 3 = 0.5222 ± 0.09894. P = 0.0272 by one-way ANOVA, post hoc Tukey’s test, * P = 0.0266. For Stage 1 vs Stage 2: # P = 0.0306 by unpaired Student’s t-test. n = 3 cells per stage. Right panel: Nonlinear fit histogram compares growth lifetime of EB3 tracks (in sec) in soma of stage 1, stage 2 and stage 3 cells. Inset shows growth half lifetime (τ) obtained from all the three stages. Mean ± SEM values of growth half lifetime (τ) obtained from stage 1 = 3.617 ± 0.4498, stage 2 = 2.616 ± 0.4468, stage 3 = 5.489 ± 0.7192. * P =, by unpaired Student’s t-test. n = 3 cells per stage. **c.** Linear regression histograms of EB3 track growth speed (µm/sec), displacement of EB3 comets (in µm) per each frame (= 2sec) and growth lifetime of EB3 tracks (in sec) in the soma are plotted against the distance from the MTOC, separately, for stage 1, stage 2 and stage 3 groups. Different shades of black and gray lines indicate linear regression for individual cells analyzed per stage, whereas the red line is the linear regression for all the cells per stage, for which Pearson correlation coefficient (r value) and P value of significance are included in the respective graphs.

Close analysis of our cells shows that at stage 1 the somatic microtubule dynamics vary depending on their distance with the MTOC position. Thus, the speed and length of growing microtubules are augmented near the MTOC compared to the soma periphery (Figure 2c and Supp Figure 1, 2). Concerning the growth lifetime of those microtubules, we detected a reduced lifetime close to the MTOC (Figure 2c and Supp Figure 3c). Once neurons formed neurites, we did not register somatic regional changes regarding microtubule dynamics (Figure 2c and Supp Figure 1-3c). Overall, our data show local differences in microtubule remodeling at the soma of stage 1 neurons. These differences suggest a radial organization of stable microtubules in the soma that might contribute later with the axon formation.

### Pharmacological manipulation of the cytoskeleton unmasks the relevance of radial organization of stable microtubules

Given our initial observations, we decided to test whether the microtubule remodeling in the soma influence the extending axon. We, therefore, hypothesized that if the F-actin cytoskeleton of stage 1 neurons is disrupted, the formation of more than one axon would be precluded due to the lack of stable microtubules that could support the extension of multiple axons. F-actin disruption using cytochalasin D (CytoD) is a well-known strategy that challenges neuronal polarity and produces neurons with multiple axons ^7^. However, it was never tested if the multipolarity induced by CytoD is stage dependent. To this end, we treated hippocampal neurons with 2µM CytoD right after plating (0hr) and around 30hr after plating for 2 days. At 30hr after plating most of the untreated cells were stage 2 neurons (Supp Figure 4b). Importantly, we found that CytoD treatment of cells at 0hr did not produce multiple axons compared to control DMSO-treated cells (Supp Figure 4a, c). However, neurons treated 30hr after plating increased the proportion of neurons producing multiple axons compared with the DMSO-treated neurons (Supp Figure 4a, c). To test if the lack of several axons at 0hr is due to the deficiency of acetylated microtubules/stable microtubules, we treated cells with the microtubule-stabilizing drug Taxol at 5nM concentration for 2 days, which was shown to produce several axons ^8^. According to our hypothesis, Taxol treatment increased the number of cells with multiple axons at both time points, 0 and 30hr, compared with control DMSO-treated neurons (Supp Figure 4a, c). Furthermore, we left our treated neurons growing for 7 days to further corroborate the axonal identity with AnkG immunostaining (which labels the axon initial segment). We found that cells treated at 0hr with CytoD did not increase the proportion of AnkG positive processes compared with DMSO-treated neurons (Supp Figure 5a-c). Conversely, neurons treated with Taxol at time 0hr produced several AnkG positive processes (Supp Figure 5a-c). These results confirm that the lack of stable microtubules at stage 1 is the limiting factor to produce multiple axons in the absence of an organized F-actin cytoskeleton.

Our plated neurons are dissociated from the tissue when they were already differentiating; thus, it is possible that our results could be affected by the dedifferentiation process that our cells were subjected to. Therefore, we decided to treat early born neurons and neurons that were already differentiating *in situ*. To this end, we *in utero* electroporated mouse cortices at embryonic day 13 (E13) and at E15 with Venus plasmid. Cortices were harvested and transfected neurons were dissociated and plated at E17. Neurons transfected at E13 were located in the cortical plate (CP) of the developing cortex; hence, they were migrating and quite advance in their differentiation process (Supp Figure 6b). Conversely, when cortices were transfected at E15 early born neurons were in the lower intermediate zone (IZ) initiating their differentiating program. Independently of the *in situ* developmental stages, we found that cortical cells treated with CytoD at time 0hr do not form multiple axons as we documented with the hippocampal neurons (Supp Figure 6a). When cultured neurons were treated with Taxol, however, they increased the proportion of multiple axons (Supp Figure 6a).

Furthermore, we performed a long-term time-lapse analysis of stage 1 and 2 hippocampal neurons treated with CytoD and Taxol. We corroborated that stage 1 neurons treated with CytoD did not produce multiple axons as the Taxol treatment does. (Figure 3a, b; Videos 5-10). Finally, using STED microscopy analysis we verified that cells treated with CytoD right after plating have fewer somatic acetylated microtubules compared with cells treated with Taxol or the ones treated with CytoD 30hr after plating (Figure 3c-e). Overall, our results suggest that the number of somatic acetylated microtubules drives axon formation.

**Figure 3.**
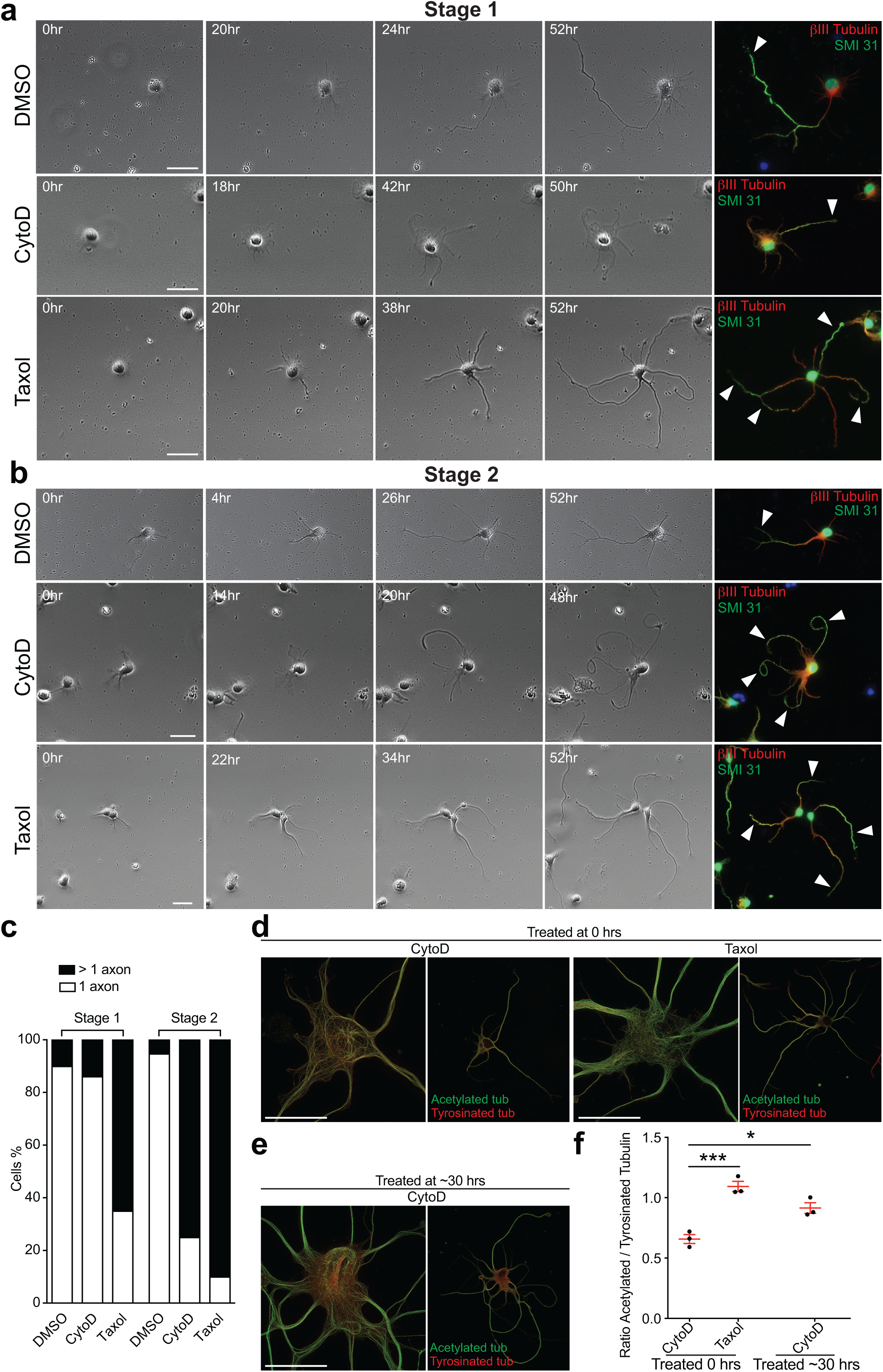
Contrary to microtubule stabilization (by Taxol), F-actin disruption (by cytochalasin D) induced multipolarity is dependent on the developmental stage of the neurons. **a-b**. Stage 1 (in a) and stage 2 (in b) primary rat hippocampal neurons followed before (0h) and during the course of 2µM cytochalasin D or 5nM Taxol compared to DMSO treatment for 52 hours, phase-contrast images of the cells with timestamps indicating the time elapsed from the start of treatment. The cells are PFA fixed for post hoc β (shown in green) immunostainings to confirm the axonal (indicated by white arrowheads) identity of the neurites. Scale bar: 10 µm. **c.** Quantifications show the percentage of stage 1 and stage 2 neurons treated with DMSO or 2µM cytochalasin D or 5nM Taxol for 52 hours differentiated to have 1 axon or more than 1 axon. DMSO treated Stage 1 cells (n = 40 cells) with 1 axon = 90 %, more than 1 axon = 10 %, CytoD treated Stage 1 cells (n = 36 cells) with 1 axon = 86.11 %, more than 1 axon = 13.89 %, Taxol treated Stage 1 cells (n = 20 cells) with 1 axon = 35 %, more than 1 axon = 65 %. DMSO treated Stage 2 cells (n = 19 cells) with 1 axon = 94.74 %, more than 1 axon = 5.26 %, CytoD treated Stage 2 cells (n = 20 cells) with 1 axon = 25 %, more than 1 axon = 75 %, Taxol treated Stage 2 cells (n = 10 cells) with 1 axon = 10 %, more than 1 axon = 90 %. **d.** STED images of acetylated and tyrosinated tubulin immunostained hippocampal neurons treated 2µM cytochalasin D immediately (0hrs) or ∼ 30 hrs after plating and 5nM Taxol immediately after plating and Inset: Centrosome labelled by Pericentrin antibody staining. White arrowheads point enrichment of tubulin acetylation in the axon. Scale bar: 10 µm. **e.** Quantifications compare ratio of acetylated to tyrosinated tubulin signal in the soma of neurons treated 2µM cytochalasin D immediately (0hrs) or ∼ 30 hrs after plating and 5nM Taxol immediately (0hrs) after plating. Mean ± SEM values of CytoD 0 hrs treated cells = 0.6572 ± 0.0370, Taxol 0hr treated cells = 1.09 ± 0.0424 and CytoD 30 hr treated cells = 0.914 ± 0.0443. P = 0.0009 by one-way ANOVA, post hoc Tukey’s test, *** P = 0.0007, * P = 0.0109. n = 3 cells per each group.

### Centriolar protein Cep120 modulates microtubules stability and axon formation

To understand mechanistically how the centrosome regulates microtubules dynamics and axon formation, we decided to interfere with the expression of Cep120, a centriolar protein, which was shown to affect microtubule stability ^4, 19, 20, 29^. Cep120 has been previously shown to control the size of the astral microtubule structure, which couples the centrosome and the nucleus in neuronal progenitors ^29^. In addition, it has been shown that Cep120 controls microtubule stability in developing neurons ^4^. Moreover, Cep120 modulates cilia formation and lack of Cep120 impairs centriole maturation, cilia elongation and microtubules acetylation on the cilia ^19, 20^. Cep120 is, therefore, a suitable candidate molecule to evaluate whether the centrosomal-dependent organization of microtubules supports axon formation. To investigate the effects of Cep120 down-regulation in early neuronal development, we used Cep120 shRNA construct for specifically silencing Cep120 expression in cortical neurons ^4^. To this end, we introduced Cep120 shRNA or control shRNA plasmids together with tDimer expressing plasmid in mice brain cortices at embryonic day 15 (E15) and isolated cortical neurons at E17. In parallel, Cep120 was over-expressed using Cep120-GFP together with tDimer expressing plasmid. Neurons were cultured for an additional 48-72hr fixed and prepared for immunostaining to assess number of neurites and axons, length of neurites, and content of somatic acetylated tubulin. Our results show that Cep120 downregulation decreased the acetylated tubulin content in the soma compared to the control-transfected neurons (Figure 4a, b). On the contrary, Cep120 overexpression increased the acetylated tubulin in the soma compared to the controls (Figure 4a, b). Importantly, we found that Cep120 overexpression produced neurons with more than one axon and overall, more complex neurons with increased length of neurites per cell, compared to control neurons (Figure 4c-f). Conversely, Cep120 downregulation decreased the complexity of the neurons with less axon per cell, fewer neurites, and reduced neurite length (Figure 4c-f). Taken together these experiments point out an important role of Cep120 in axon formation and microtubules acetylation as previously shown for the cilia formation ^19, 20^.

**Figure 4.**
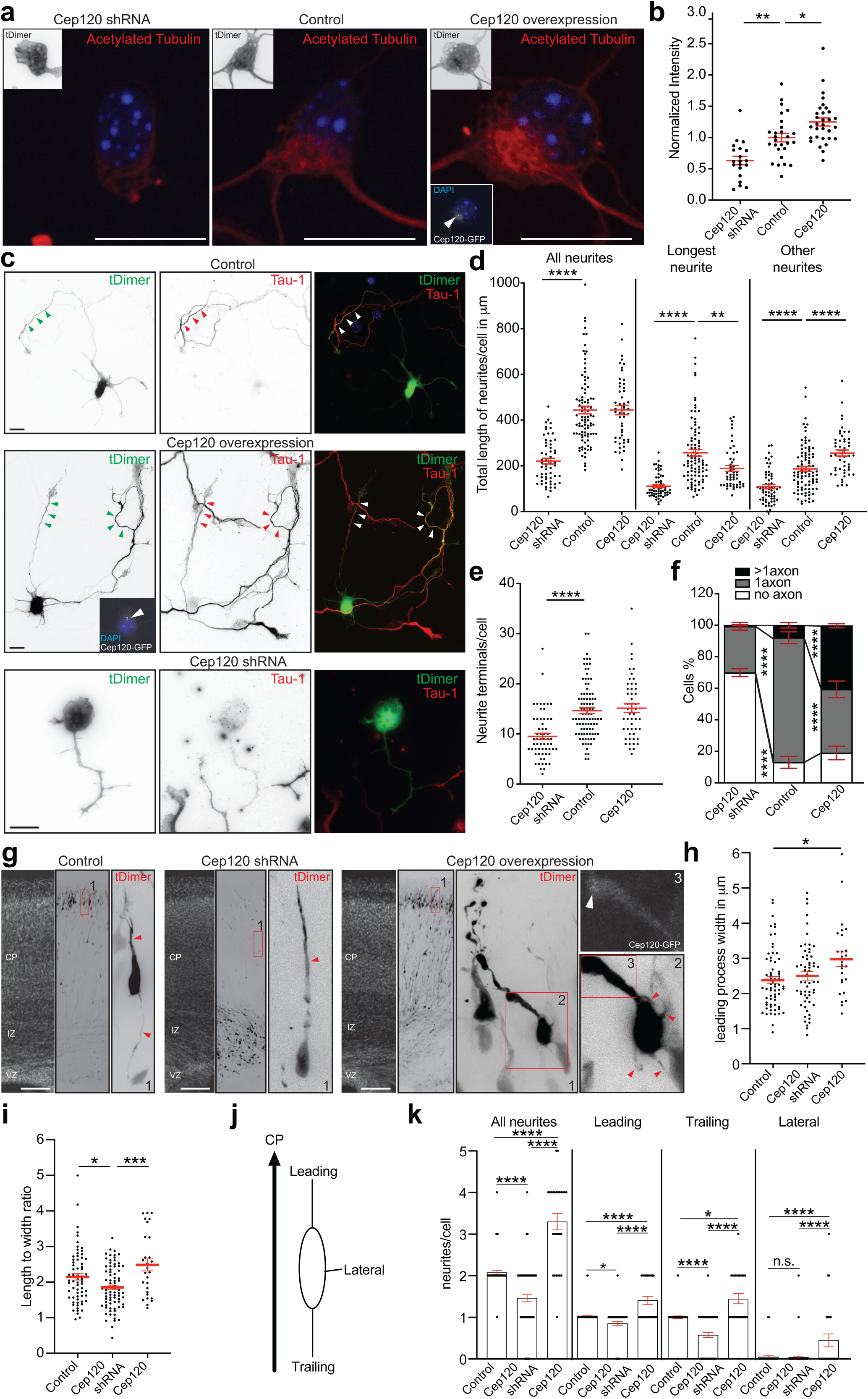
Cep120 knockdown and overexpression, through microtubule acetylation, bidirectionally regulates axon formation. **a.** Confocal maximum projection images of mouse cortical neurons co-transfected at E15 via IUE with tDimer and Cep120 shRNA, or control or Cep120-GFP (indicated by arrowhead in the inset) cultured at E17 for 48 hrs. immunostained with acetylated tubulin antibody. Scale bar: 10 µm **b.** Quantifications compare normalized acetylated tubulin intensities in the soma of neurons expressing Cep120 shRNA (n = 20), control (n = 27) and Cep120-GFP (n = 31) as shown in **a**. Mean ± SEM values of Cep120 shRNA cells = 0.6322 ± 0.0678, control cells = 1.000 ± 0.0676 and Cep120-GFP cells = 1.249 ± 0.0641. P < 0.0001 by one-way ANOVA, post hoc Tukey’s test, **** P < 0.0001, ** P = 0.0014. **c.** Epifluorescence images of mouse cortical neurons co-transfected via IUE at E15 with tDimer and Cep120 shRNA, or control or Cep120-GFP (indicated by arrowhead in the inset) cultured at E17 for 48 or 72 hrs. immunostained with Tau-1 antibody to confirm the axonal identity of the neurites. Scale bar: 10 µm **d.** Neurite length quantification in neurons (in µm) expressing Cep120 shRNA (n = 60), control (n = 93) and Cep120-GFP (n = 51) as shown in **c**. Mean ± SEM values for length of all neurites in, Cep120 shRNA cells = 220.4 ± 12.14, control neurons = 441.1 ± 16.62, Cep120-GFP cells = 444.4 ± 20.25. Mean ± SEM values for length of longest neurite in Cep120 shRNA cells = 111.4 ± 6.56, control neurons = 257.1 ± 14.65, Cep120-GFP cells = 188.2 ± 12.67. Mean ± SEM values for length of other neurites in Cep120 shRNA cells = 109 ± 9.062, control neurons = 186.9 ± 10.46, Cep120-GFP cells = 256.2 ± 13.36. P < 0.0001 by one-way ANOVA, post hoc Tukey’s test, **** P < 0.0001, ** P = 0.011. **e.** Neurite terminals quantification in neurons expressing Cep120 shRNA (n = 60), control (n = 93) and Cep120-GFP (n = 51) as shown in **c**. Mean ± SEM values for neurite terminal per cell in Cep120 shRNA cells = 9.517 ± 0.6127, control neurons = 14.66 ± 0.5752, Cep120-GFP cells = 15.16 ± 0.8643. P < 0.0001 by one-way ANOVA test, post hoc Tukey’s test, **** P < 0.0001. **f.** Quantifications show the percentage of Cep120 shRNA (n = 258), control (n = 364) and Cep120-GFP (n = 172) neurons, as shown in **c**, differentiated to have no axon, 1 axon and more than 1 axon. Mean ± SEM values for percentage of Cep120 shRNA cells with no axon = 69.90 ± 2,450, 1 axon = 29.43 ± 2.404, more than 1 axon = 0.7000 ± 0.3606; percentage of control cells with no axon = 13.01 ± 3.762, 1 axon = 79.10 ± 3.785, more than 1 axon = 7.892 ± 1.677; percentage of Cep120-GFP cells with no axon = 19.01 ± 4.226, 1 axon = 40.30 ± 5.231, more than 1 axon = 40.44 ± 1.279. α = 0.05 by two-way ANOVA, post hoc Tukey’s test, **** P < 0.0001. Data shown in **b**, **d, e** and **f** is obtained from cortical cultures of IUE mouse embryos from at least 2 different mothers. **g.** E19 mouse cortical slices co-transfected via IUE at E15 with tDimer and Cep120 shRNA, or control or Cep120-GFP (indicated by white arrowhead in the inset 3). Red insets (labelled 1): zoomed view of migrating neurons in the cortical plate from each condition are shown. Leading and trailing processes are indicated by red arrow heads. Cep120 down regulation precluded trailing process, whereas Cep120-GFP overexpression increased the number of neurites as indicated by the red arrow heads in the inset 2. Scale bar: 200 µm. **h.** Quantifications show leading process width (in µm) of neurons in the cortical plate expressing Cep120 shRNA (n = 59), control (n = 67) and Cep120-GFP (n = 27) as shown in **g**. Mean ± SEM values of leading process width in Cep120 shRNA cells = 2.501 ± 0.1199, controls = 2.385 ± 0.1031, Cep120-GFP cells = 2.976 ± 0.2061. P = 0.0204 by one-way ANOVA, post hoc Tukey’s test, * P = 0.0155. **i.** Quantifications show length to width ratio of the soma of neurons in the cortical plate expressing Cep120 shRNA, control and Cep120-GFP expressing neurons as shown in **g**. Mean ± SEM values of length to width ratio of soma in Cep120 shRNA cells = 1.852 ± 0.0664, control cells = 2.148 ± 0.0927, Cep120-GFP cells = 2.485 ± 0.1683. P = 0.0002 by one-way ANOVA, post hoc Tukey’s test, *** P = 0.0002, * P = 0.0344. **j.** Schematic illustrating categorization of apical, basal and lateral neurites used for analyzing data in **k**. **k.** Neurite number quantifications of neurons in the cortical plate expressing Cep120 shRNA, control and Cep120-GFP expressing neurons as shown in **g**. Mean ± SEM values for all neurites in Cep120 shRNA cells = 1.463 ± 0.0906, control cells = 2.077 ± 0.0505, Cep120-GFP cells = 3.296 ± 0.1984. Mean ± SEM values for apical neurites in Cep120 shRNA cells = 0.8500 ± 0.0440, control neurons = 1.031 ± 0.0216, Cep120-GFP cells = 1.407 ± 0.0964. Mean ± SEM values for basal neurites in control neurons = 1.000 ± 0.0310, Cep120 shRNA cells = 0.5750 ± 0.0610, Cep120-GFP cells = 1.444 ± 0.1233. Mean ± SEM values for lateral neurites in Cep120 shRNA cells = 0.0375 ± 0.0278, control cells = 0.0461 ± 0.0262, Cep120-GFP cells = 0.4444 ± 0.1541. P < 0.0001 by Kruskal-Wallis ANOVA test, post hoc Dunn’s test, **** P < 0.0001. * P = 0.0112, n.s. = not significant. Data shown in **h**, **i** and **k** is obtained from three E19 mouse brains slices from two Cep120-GFP and three control and three Cep120 shRNA conditions.

To further investigate the role of Cep120 on neuronal differentiation *in vivo*, we decided to overexpress or downregulate Cep120 in the developing cortex. In previous work, we found that specific downregulation of Cep120 in the developing cortex precludes axon formation and impairs neuronal migration ^4^. It was not investigated, however, whether the Cep120 overexpression affects neuronal development *in vivo*. To test this, we *in utero* electroporated cortices with Cep120 shRNA or Cep120 overexpressing plasmids together with tDimer plasmid at E15 and brains were harvested at E19. We specifically analyzed the migrating neurons in the cortical plate (CP) when they already should have formed an axon. Our results show that Cep120 downregulation leads to alterations of the cell body morphology with a reduction of the length to width ratio compared with control transfected neurons (Figure 4g, i). In addition, we detected a diminution of bipolar neurons at the CP without trailing process or future axon compared with control neurons (Figure 4g, j, k). In few neurons, we also observed that Cep120 downregulation precluded the formation of a leading process (Figure 4k). On the contrary, Cep120 overexpression produced migrating neurons in the CP with increased width of the leading process (Figure 4g, h). Some Cep120-transfected neurons bear more than one leading process or trailing process and overall, the neurons overexpressing Cep120 have processes emerging from the cell body, which we called lateral processes, that the control transfected neurons do not have (Figure 4g, j, k). Altogether these results demonstrate that Cep120 downregulation or overexpression has a bidirectional effect on the morphological complexity of migrating neurons in the CP.

### Lack of Cep120 affects the landscape of proteins regulating microtubule dynamics

In order to test more directly the effect of Cep120 downregulation, we decided to measure the proteome in the absence of Cep120 *in vivo*. To this end, we *in utero* electroporated cortices at E15 with Cep120 shRNA + pNeuroD-GFP (which is expressed exclusively in neurons ^4^) or control plasmid together with pNeuroD-GFP, at E19 brains were harvested and GFP positive cells were FAC sorted. Afterwards, cells were prepared for differential quantitative proteomics and bioinformatics analyses (Figure 5a). Our results show that from the 1093 quantifiable proteins 186 were significantly altered after Cep120 downregulation (Figure 5b). Interestingly, among these several proteins associated with microtubule dynamics were downregulated in the absence of Cep120, such as Map2, Tau, EB1, Dcx, CRMP2, MAP1b, and Tubb2b among others (^30, 31^; Figure 5c, d; Supp. Figure 7). Finally, gene ontology (GO) analysis supported this general impression since several categories linked to axonal elongation and formation were enriched in the control sample vs. neurons missing Cep120. In detail, gene set enrichment analysis (GSEA) across all identified proteins across the two samples indicates that categories such as axon development (*Biological process*), site of polarized growth, microtubule associated complex, and microtubule (*Cellular component*) were significantly enriched in control samples *vs.* neurons missing Cep120 (Figure 5e). Altogether, these results suggest substantial changes to the proteome landscape of neurons without Cep120, which might affect microtubules dynamics and axon formation/extension.

**Figure 5.**
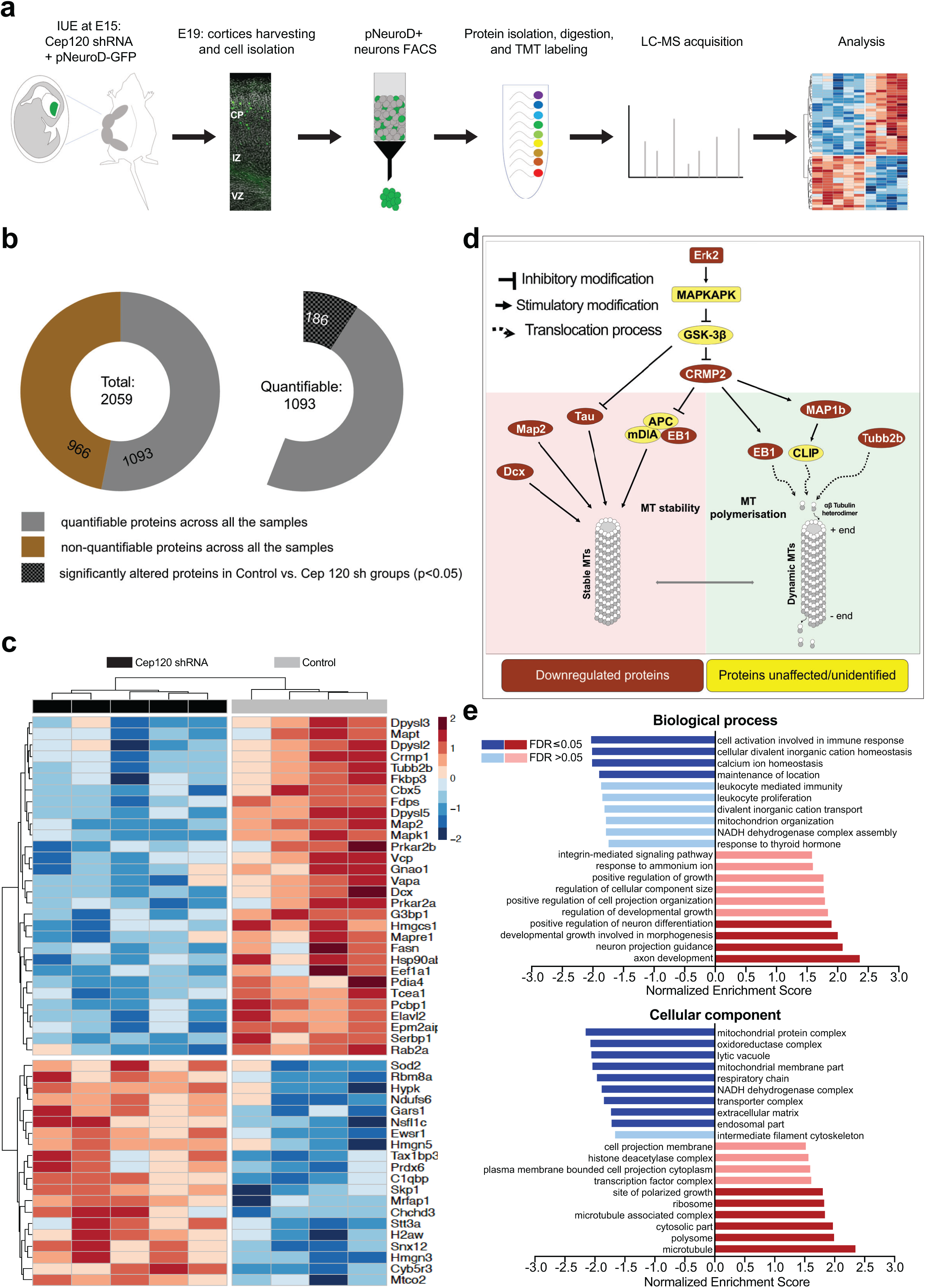
Cep120 knockdown downregulates key proteins related to microtubule stability and dynamics - revealed by differential quantitative mass spectrometric proteome analysis of migrating cortical neurons. **a.** Schematic drawing of different steps involved in the proteome analysis performed on migrating cortical neurons at E19 transfected via IUE at E15 with control or Cep120 shRNA along with pNeuroD-GFP plasmid. **b.** Illustration of total number of quantifiable (966) and non-quantifiable (1093) proteins identified in the proteome analysis across all the samples: n = 4 for controls and n = 5 for Cep120 shRNA condition. Out of the 1093 proteins quantified, 186 are significantly altered in control vs Cep120sh groups, with P < 0.05. (See Supplementary excel file Rawdata.xlsx for details). **c.** Heatmap of 50 significantly altered proteins in the samples from control vs Cep120 knockdown conditions with the lowest significant p-value (< 0.05). The dendrogram represents the hierarchical clustering with a distance score based on Pearson-correlation coefficients (z-score -2 to 2) and a complete linkage clustering as cluster agglomeration criteria ((dis-)similarity between Cep120 shRNA and control groups). **d.** Pathway illustrates key proteins that are downregulated upon Cep120 knockdown, obtained from the top 50 altered proteins list with the lowest significant p-value (< 0.05) shown in **c**. Alternate names of some of the affected proteins listed in the pathway: Erk2 = MAPK1, CRMP2 = Dpysl2, Tau = MAPT, EB1 = Mapre1. Pathway modified and reproduced from: Wiki pathways: https://www.wikipathways.org/index.php/Pathway:WP2038; Cell Signaling Technology Pathways: https://www.cellsignal.de/contents/science-cst-pathways-adhesion-ecm-cytoskeleton/regulation-of-microtubule-dynamics/pathways-micro **e.** Gene set enrichment analysis (GSEA) analysis of control samples *vs.* Cep120 shRNA resuming the results the two non-redundant functional ontology databases: biological process and cellular component. The analysis was performed with WebGestalt for the 1093 proteins quantified in both sample groups. (See Supplementary excel file GSEA.xlsx for details). FDR: false discover rate

### Cep120 affects microtubule dynamics

To test if Cep120 manipulation affects microtubule dynamics, we decided to monitor growing microtubules by time-lapse microscopy using tdTomato-EB3. Cortical neurons were *in utero* electroporated at E15 with Cep120 shRNA + tdTomato-EB3 or Cep120-GFP + tdTomato-EB3. At E17 transfected neurons were isolated from the cortices, cultured, and prepared for live imaging. 24hr after plating stage 2 neurons were imaged for 5 minutes with a frame rate of 2 seconds (Videos 11-13). Our results show that the median growth speed of microtubules is increased significantly after Cep120 overexpression compared with control transfected neurons (Figure 6a, b). Cep120 downregulation showed a tendency (p = 0.060, by unpaired t test) to have a decreased median speed per cell compared with control transfected neurons (Figure 6a, b). However, when speed of EB3 tracks from all the cells were plotted, we found significant reduction of the growth speed after Cep120 downregulation (Supp. Figure 8a, b). Likewise, the characteristic growth per frame (λ) shows higher EB3 comet displacement when Cep120 is overexpressed (Figure 6a, b) and a tendency of reduction (p = 0.065, by unpaired t test) in neurons lacking Cep120. Considering the median displacement of all EB3 comets recorded, we found a significant reduction in the displacement after Cep120 downregulation (Supp. Figure 8c, d). Finally, we analyzed the characteristic lifetime (τ) per cell and no significant differences were detected between control transfected neurons and Cep120-overexpressing or Cep120-downregulated neurons (Figure 6a, b). However, the median of the lifetime of all the recorded tracks per cell showed differences between the control neurons and the neurons in which Cep120 levels were manipulated: a significant increment of the median lifetime after Cep120 downregulation and a significant decrement with Cep120 overexpression (Supp. Figure 8e, f). These results highlight that overexpression of Cep120 levels lead to increased microtubule dynamics. On the contrary, Cep120 downregulation decreased microtubule dynamics.

**Figure 6.**
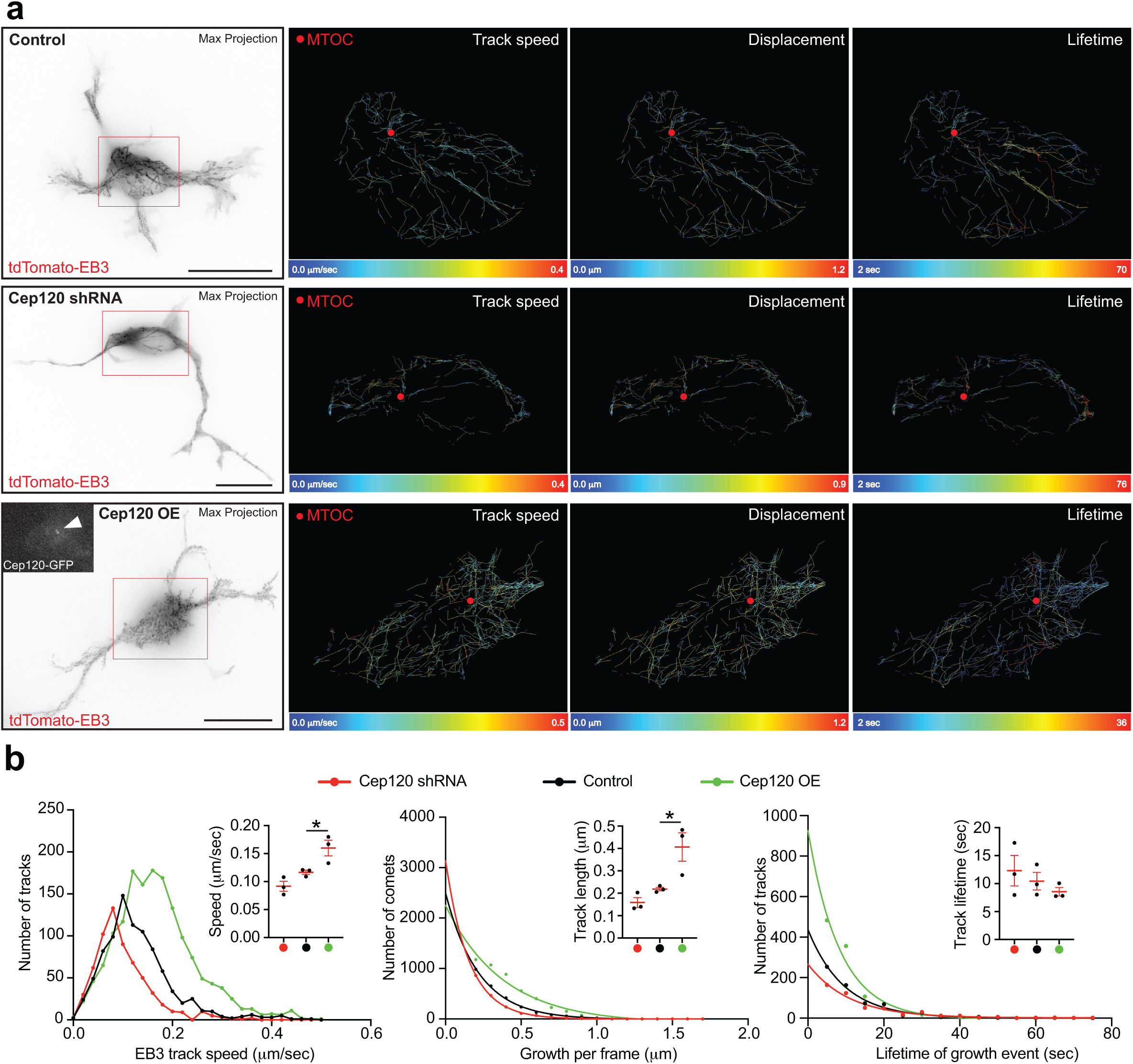
Microtubule plus-end (EB3) dynamics are modulated by Cep120. **a.** Images showing maximum intensity projections (black on white images) from a 5 min time-lapse (1 frame every 2 seconds) of stage 2 mouse cortical neurons co-transfected (via IUE at E15) with EB3-tdTomato and control or Cep120 shRNA or Cep120-GFP (indicated by arrowhead in the inset). Speed of EB3 tracks, displacement (per frame) and lifetime of EB3 tracks (analysed by TrackMate ImageJ plugin) in the soma from the respective cells were shown as heatmaps. Red dot in the heatmaps indicate MTOC. Scale bar: 10 µm. **b.** Left panel: Frequency distribution histogram compares Speed of EB3 tracks (µm/sec) in the soma of Cep120 shRNA, control, and Cep120-GFP expressing neurons. Inset shows values of median speed (µm/sec) obtained from all the three groups. Mean ± SEM values of median speed in the soma of Cep120 shRNA cells = 0.0920 ± 0.0087, control cells = 0.1167 ± 0.0038, Cep120-GFP cells = 0.1600 ± 0.0140. P = 0.0074 by one-way ANOVA, post hoc Tukey’s test, * P = 0.0462. n = 3 cells per group. Middle panel: Nonlinear fit histogram compares displacement of EB3 (in µm) per each frame (2sec) in the soma of Cep120 shRNA, control, and Cep120-GFP expressing neurons. Inset shows characteristic growth (λ) per frame, which is the half values of displacement per frame obtained from all the three groups. Mean ± SEM values of the characteristic growth (λ) per each frame in the soma of Cep120 shRNA cells = 0.1585 ± 0.0224, control cells = 0.2185 ± 0.0080, Cep120-GFP cells = 0.4068 ± 0.0633. P = 0.0098 by one-way ANOVA, post hoc Tukey’s test, * P = 0.0330. n = 3 cells per group. Right panel: Nonlinear fit histogram compares growth lifetime of EB3 tracks (in sec) in the soma of Cep120 shRNA, control, and Cep120-GFP expressing neurons. Inset shows growth half lifetime (τ) obtained from all the three groups. Mean ± SEM values of growth half lifetime (τ) in the soma of Cep120 shRNA cells = 11.13 ± 2.090, control cells = 10.41 ± 1.555, Cep120-GFP cells = 8.513 ± 0.7535. P = 0.5743 by one-way ANOVA, post hoc Tukey’s test, n.s. = not significant. n = 3 cells per group.

### Cep120 restores microtubules acetylation and promotes axon formation

Finally, we decided to test whether Cep120 could restore microtubule dynamics, and thus axon formation, when microtubule dynamics are affected. Previously, it was shown that Cep120 in coordination with the transforming acid coiled-coil protein 3 (TACC3), which is a microtubule plus end protein ^32^, promote the elongation of the microtubule aster in neuronal progenitors ^29^. Moreover, TACC3 promotes axon elongation and regulates microtubule plus end dynamics ^32–34^. Therefore, TACC3 is a suitable candidate to challenge neuronal microtubule dynamics and test the effect of Cep120 under those constraints. We used a TACC3 inhibitor (SPL-B) that selectively inhibits the nucleation of centrosome microtubules ^35^. We found that SPL-B decreased the somatic content of acetylated microtubules compared with control treated neurons (Figure 7a, b). However, overexpression of Cep120 increased the content of somatic acetylated microtubules in the presence of SPL-B (Figure 7a, b). Moreover, SPL-B treated cells decreased the length and number of neurites per cell and preclude axon formation (Figure 7c-f). Similar results were obtained when TACC3 was downregulated with already published and validated shRNA sequence (^33^; Supp. Figure 9). Importantly, Cep120 overexpression overcomes partially these SPL-B-dependent deficits and neurons initiated to form neurites and to extend an axon (Figure 7c-f). Altogether, these results demonstrate that increased levels of Cep120 are able to overcome insufficiencies with microtubule stability that preclude neuronal differentiation and axon formation.

**Figure 7.**
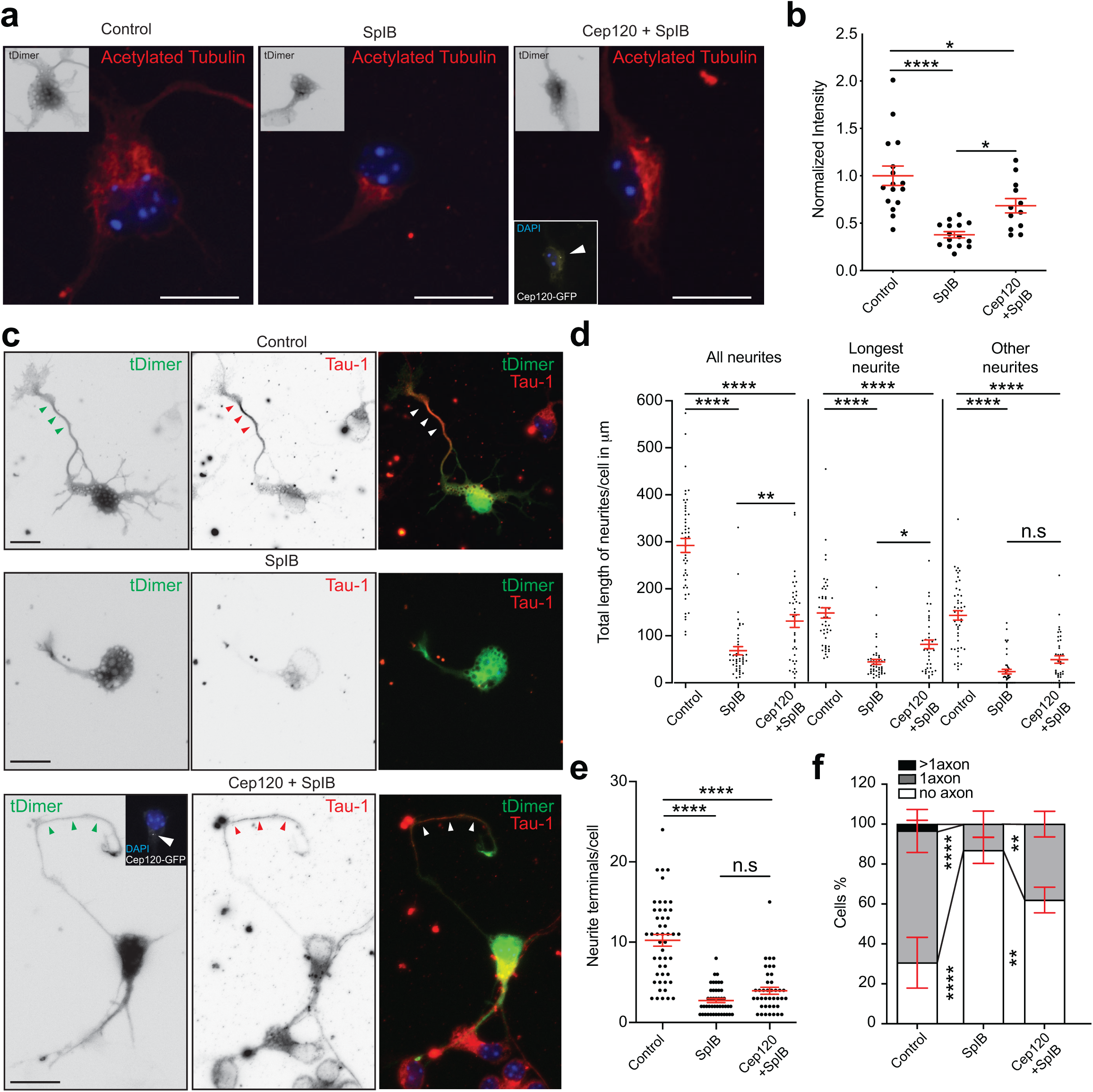
Cep120 rescues SPL-B induced axon loss phenotype, by promoting microtubule acetylation. **a.** Confocal maximum projection images of mouse cortical neurons co-transfected via IUE at E15 with tDimer alone or together Cep120-GFP (indicated by arrowhead in the inset) cultured at E17 were treated immediately after plating with 0.25µg/ml Spindlactone B (SPL-B, a TACC3 inhibitor) for 48 hrs. immunostained with acetylated tubulin antibody. Scale bar: 10 µm. **b.** Quantifications compare normalized acetylated tubulin intensities in the soma of control (n = 16), SPL-B-treated cells (n = 15) and SPL-B treated Cep120-GFP neurons (n = 12) as shown in **a**. Mean ± SEM values for the soma of control cells = 1.000 ± 0.1029, SPL-B cells = 0.3778 ± 0.0325, and Cep120-GFP+SPL-B cells = 0.6836 ± 0.0758. P < 0.0001 by one-way ANOVA, post hoc Tukey’s test, **** P < 0.0001, * P < 0.05. **c.** Epifluorescence images of mouse cortical neurons co-transfected via IUE at E15 with tDimer alone or together Cep120-GFP (indicated by arrowhead in the inset) cultured at E17 were treated immediately after plating with 0.25µg/ml SPL-B for 48 hrs. immunostained with Tau-1 antibody. Scale bar: 10 µm. **d.** Neurite length quantification (in µm) of control (n = 48), SPL-B treated (n = 45) and SPL-B treated Cep120-GFP (n = 40) neurons as shown in **c**. Mean ± SEM values for length of all neurites of control cells = 292.2 ± 15.06, SPL-B cells = 68.20 ± 8.768, and Cep120-GFP + SPL-B cells = 131.4 ± 13.60. Mean ± SEM values for length of longest neurite in control cells = 148.7 ± 10.96, SPL-B cells = 44.57 ± 5.252, Cep120-GFP cells = 81.89 ± 9.284. Mean ± SEM values for length of other neurites in control cells = 143.5 ± 10.01, SPL-B cells = 23.63 ± 4.889, and Cep120-GFP + SPL-B cells = 49.47 ± 7.975. P < 0.0001 by one-way ANOVA, post hoc Tukey’s test, **** P < 0.0001, ** P = 0.0026, * P = 0.0126, n.s. = not significant. **e.** Neurite terminals quantification in control (n = 48), SPL-B -treated cells (n = 45) and SPL-B treated Cep120-GFP neurons (n = 40) as shown in **c**. Mean ± SEM values for neurite terminal per cell for controls = 10.23 ± 0.7313, control + SPL-B cells = 2.733 ± 0.2551, and Cep120-GFP + SPL-B cells = 3.950 ± 0.4370. P < 0.0001 by one-way ANOVA, post hoc Tukey’s test, **** P < 0.0001, n.s. = not significant. **f.** Quantifications show the percentage of control (n = 123 cells), SPL-B-treated (n = 106 cells) and SPL-B treated Cep120-GFP (n = 56 cells) neurons, as shown in **c**, differentiated to have no axon, 1 axon and more than 1 axon. Mean ± SEM values for percentage of control cells with no axon = 30.57 ± 12.71, 1 axon = 66.01 ± 10.79, more than 1 axon = 3.425 ± 1.935; percentage of SPL-B cells with no axon = 86.83 ± 6.500, 1 axon = 13.17 ± 6.500, more than 1 axon = 0.000 ± 0.000; percentage of SPL-B treated with Cep120-GFP cells with no axon = 61.99 ± 6.430, 1 axon = 38.01 ± 6.430, more than 1 axon = 0.000 ± 0.000. α = 0.05 two-way ANOVA, post hoc Tukey’s test, **** P < 0.0001, ** P = 0.0013. Data shown in **b**, **d**, **e** and **f** is obtained from cortical cultures of IUE mouse embryos from at least 2 different mothers.

## Discussion

### Polarity before polarization

Tubulin post-translation modifications (PTMs), and specifically acetylation, detyrosination and other polymodifications have crucial roles in the assembly, maintenance and function of complex and stable microtubule-based organelles that form the core components of the centrosome such as the centrioles, basal bodies (the protein structure at the base of a cilium or flagellum) and axonemes (the central strand of a cilium or flagellum) (reviewed in ^36^). In addition, various studies point out the presence of stable microtubules in axons - determined either by measuring acetylated ^8^ or detyrosinated microtubules ^37^. Overall, it seems that axons contain a higher percentage of stable microtubules compared to their dendritic counterparts (reviewed in ^38^). Yet, it is not completely clear the exact mechanisms by which differential microtubule stability is achieved in axons versus other minor neurites (future dendrites).

Our results demonstrate that before axon extension, the soma contains acetylated microtubules preferentially surrounding the centrosome. Once neurites are formed the acetylated microtubules penetrate them to be eventually enriched in the growing axon. This initial somatic microtubule organization might set the conditions to break the symmetry once axonal fate is defined.

Supporting this view, we uncovered differential effects of CytoD, which is known to induce multiple axons by causing F-actin disruption, on stage 1 and stage 2 cells. Unlike Taxol (that causes microtubule stability), CytoD did not induce multiple axons in the less developed stage 1 cells, with relatively fewer stable microtubules in the soma. Accordingly, we found that Cep120, a centriolar protein that promotes microtubules stability ^19–21^, affects the axon formation. Importantly, we could show that the levels of Cep120 modulate axon formation in a bidirectional manner. Consequently, lack of Cep120 precluded axon formation, and excess of Cep120 produced neurons with multiple axons. Moreover, we found that lack of Cep120 affects the content of proteins associated with microtubule dynamics and stability, such as EB1, CRMP2, and Tau. Although the exact molecular mechanisms by which Cep120 orchestrates the levels of these proteins remains for the moment undetermined, our data suggest that the centrosome might have a central role in initiating neuronal differentiation by regulating microtubule dynamics.

### Microtubule dynamics versus stability

We found in early differentiating neurons (stage 1) regional somatic microtubule dynamics with increased growth speed and length near the MTOC and decreased lifetime of the EB3 trajectories. Thus, suggesting highly dynamic microtubules near the MTOC. In fixed samples, still, we detected preferentially acetylated microtubules suggesting stable microtubules near the centrosome. Of note, a long-standing question in cell biology is whether acetylated tubulin alters microtubule polymerization dynamics. In this regard, it was shown that promoting pharmacologically tubulin acetylation did not alter microtubule dynamics (e.g., growth rate and shortening rate). Nevertheless, those acetylated microtubules were more resistant to the depolymerization effect of nocodazole ^28^. Consequently, proposing that stable microtubules do not mean fewer dynamic microtubules.

At later stages, we could not uncover somatic regional differences regarding speed, length, or lifetime of growing microtubules. However, at stage 2, previous to axon formation, we detected an overall increment of somatic EB3 speed and track length. Similarly, stage 2 cells overexpressing Cep120, which eventually formed multiple axons, showed an increment on those parameters compared with control cells. Overall, these results suggest local microtubules remodeling at stage 1, whereas the stage 2 is the transitional stage that might sustain drastic microtubule changes in the soma in order to set conditions for symmetry breakage, leading to axon elongation.

### Axon elongation

Selective translocation of the Kinesin-1 motor domain into the nascent axon was described as one of the earliest events during the axon elongation ^39^. Tubulin acetylation and detyrosination were shown to be required for Kinesin-1 motor domain translocation into the axons ^40, 41^. A later study, however, showed that acetylation of microtubules itself is not enough for sorting of Kinesin 1 into axons ^42^. Moreover, taxol-induced translocation of Kinesin 1 into supernumerary axons correlated with the enhancement of three different microtubule PTMs (acetylation, detyrosination, and glutamylation; ^42^).

Once the axon is growing, the organization of microtubule arrays shifts from centrosome-dependent to centrosome-independent in hippocampal neurons in culture. Centrosome ablation during axon elongation does not affect axon extension or regeneration ^6^. Yet, centrosome ablation in stage 2-early stage 3 neurons decreased the content of somatic growing microtubules ^43^. Furthermore, centrosome ablation in neurons developing *in vivo*, before any sign of axon development, preclude axon formation ^3^. In more developed stage 3 neurons, however, there is decentralization of centrosomal proteins, such as γ-tubulin, which in turn can organize acentrosomal microtubules in older neurons ^6^. Along these lines, Augmin and γ-TuRC are shown to be crucial for uniform plus-end out microtubule polarity in axons of matured neurons ^44^. CAMSAP2, a microtubule minus-end protein can establish and maintain non-centrosomal microtubule networks and is suggested to be crucial for axon specification and polarity of neurons *in vivo* and *in vitro* ^45^. However, recently it has been described that CAMSAP3, but not CAMSAP2, affects axon formation ^46^. CAMPSAP3 regulates microtubules stability and its absence promotes microtubules acetylation leading to the formation of multiple axons. In contrast, the lack of CAMSAP2 did not affect neuronal polarity ^46^ as suggested earlier ^45^. Nevertheless, the molecular pathways instructing the centrosome to lose its initial role during neuronal differentiation are not completely understood. In other words: how the microtubule remodeling shift from the centrosomal to an acentrosomal manner? We believe our data will pave the way to better understand the different aspects of neuronal differentiation and help to conciliate the role of the centrosome with the initial axon extension.

## Supporting information

GSEA

Rawdata

STRING

## Materials and Methods

### RNAi and fluorescent protein constructs

We previously reported the Venus, Farnesylated-GFP, pNeuroD-GFP, control shRNA, Cep 120 shRNA, and Cep 120-GFP plasmids (de anda et al 2010, Xie et al 2007). tDimer (pAAV-CAG-tDimer) was kindly provided by Thomas Oertner (ZMNH, UKE). EB3-tdTomato was a gift from Erik Dent (Addgene plasmid # 50708;http://n2t.net/addgene:50708; RID:Addgene_50708), pBrain-GFP-shTACC3 shRNA was a gift from Stephen Royle (Addgene plasmid # 59355; http://n2t.net/addgene:59355; RRID: Addgene_59355), pBrain-GFP-shGL2 was a gift from Stephen Royle (Addgene plasmid # 60004; http://n2t.net/addgene:60004; RRID:Addgene_60004).

### Animal experiments

Animal (rat and mouse) experiments were performed according to the German and European Animal Welfare Act and with the approval of local authorities of the city-state Hamburg (Behörde für Gesundheit und Verbraucherschutz, Fachbereich Veterinärwesen) and the animal care committee of the University Medical Center Hamburg-Eppendorf.

### *In utero* electroporation (IUE)

Pregnant C57BL/6 mice with E13 or E15 embryos were first administered with pre-operative analgesic, buprenorphine (0.1 mg/kg), by subcutaneous injection. After 30 min, mice were anesthetized with isoflurane (4% for induction, 2–3% for maintenance) in oxygen (0.5–0.8 l/ min for induction and maintenance). Later, uterine horns were exposed, and plasmids mixed with Fast Green (Sigma) were microinjected into the lateral ventricles of embryos. Five current pulses (50 ms pulse/950 ms interval) at 30V for E13 and 35V for E15 were delivered across the heads of embryos. After surgery, mice were kept in a warm environment and were provided with moist food containing post-operative analgesic, meloxicam (0.2–1 mg/kg), until they were euthanized for collection of the brains from the embryos. The brains were either used for cortical cultures or cortical slices or FACS.

### Mouse primary cortical cultures

Brain cortices transfected via IUE at E13 or E15 with control or Cep120 shRNA or Cep120-GFP or TACC3 shRNA or GL2 shRNA plasmids in combination with tDimer or EB3-tdTomato or Venus alone were used for cortical cultures. The concentration of shRNA (control or Cep120 shRNA or TACC3 shRNA, GL2 shRNA), Cep120 GFP plasmids injected was 2-3-fold higher than that of the tDimer plasmids or EB3-tdTomato. We used 1.5 µg/µl for shRNA (control or Cep120 shRNA or TACC3 shRNA), 1.2 µg/µl Cep120-GFP, 0.5 µg/µl of tDimer or EB3- tdTomato and 0.5 µg/µl of Venus plasmid. Two days later pregnant mice were anesthetized with CO2/O2, euthanized before taking the E17 embryos out from their uteri. Embryos were then decapitated, skulls were opened, brains were collected in petri dishes with Hibernate^TM^-E medium (Invitrogen) on ice. Hemispheres were separated, meninges were carefully stripped away, and cortices were dissected on ice. Transfected (fluorescent) cortical regions were identified and dissected on ice under a stereo microscope (Olympus SZX16) equipped with a UV light source. The isolated cortical regions were first incubated in 1x HBSS (Invitrogen) with papain and DNase (Worthington) for 10 min at 37°C neurons and then triturated. The cells were then pelleted and washed with fresh HBSS before they were plated on poly-L-lysine coated coverslips or tissue culture chambers (Sarstedt, for live imaging) in Neurobasal/B27 medium (Invitrogen), maintained in culture for 24 to 72 hrs at 37°C with 5% CO2 before use.

### Mouse cortical slices

We introduced Cep120 shRNA or control shRNA or Cep120-GFP plasmids in combination with tDimer plasmid into brain cortices at E15. We used 1.5 µg/µl for shRNA (control or Cep120 shRNA), 1.2 µg/µl of Cep120-GFP and 0.5 µg/µl of tDimer. The brains collected from E19 embryos were post-fixed in 4% paraformaldehyde (PFA) overnight at 4°C and later moved to 30% sucrose (in PBS) until they were completely sunk. Brains were then embedded in Tissue Tek OCT compound and stored at 80°C until they were sectioned to 60 µm slices using a cryostat.

### Rat primary hippocampal neuron cultures and transfections

Pregnant rats were anesthetized with CO2/O2, euthanized before taking the E18 embryos out from their uteri. Embryos were then decapitated, skulls were opened, brains were collected in petri dishes with HBSS on ice. Hemispheres were separated, meninges were carefully stripped away, and hippocampi were dissected on ice and triturated in 1xHBSS (Invitrogen) after digestion by papain and DNase (Worthington for 10 min at 37°C). Transfections were performed using the Amaxa nucleofector system following the manufacturer’s manual. The final concentration for the EB3-tdTomato or Farnesylated-GFP plasmid was 1 µg and empty pcDNA 3.1 was used to make up to 3 µg of DNA for for 5 × 10^6^ cells per each transfection mix as per the manufacturer recommendation. After electroporation, neurons were plated on poly-L-lysine coated coverslips or tissue culture chambers (Sarstedt, for live imaging) in Neurobasal/B27 medium (Invitrogen) and were maintained in culture for 24 to 72 hrs or 7-8 days at 37°C with 5% CO2 before use.

### Pharmacological treatments

Pharmacological compounds were directly added to rat hippocampal or mouse cortical neurons in culture. CytoD (Sigma) - 2µM and Taxol (Sigma) - 5nM were added either at the time of plating (at 0h) or ∼30h after plating. Spindlactone B (SPL-B; a TACC3 inhibitor; Axon Medchem) – 0.25 µg per ml was added at the time of plating (at 0h). The treated cells were PFA (4%) fixed after 48 or 72h in culture for immunostaining.

### Immunocytochemistry

Rat hippocampal or mouse cortical neurons grown on coverslips were fixed with 4% paraformaldehyde (PFA) at 37°C for 10 min and then permeabilized with 0.5% Triton X-100 for 10 min. Non-specific binding was blocked by incubation with 5% donkey serum in PBS for 60 min at RT (room temperature), followed by specific primary antibody incubation: (Tau-1 – 1:700, Acetylated Tubulin – 1:1000, SMI 31 (1:300), Ankyrin-G 1:500, βIII Tubulin – 1:2000) was added for incubation for 180 min at RT or overnight at 4°C, followed by 3 times 4 minutes PBS washes. Coverslips were incubated with respective anti-mouse or anti-rabbit Alexa Fluor 488 or 568 or 647 secondary antibodies (Invitrogen), along with Hoechst dye (1:10,000, Invitrogen) to stain for nuclei for 60 min at RT followed by three washing steps with PBS. Coverslips were mounted onto slides using Fluoromount-G® (SouthernBiotech) and were stored protected from light.

### Epifluorescence imaging

Imaging was performed on an inverted Nikon microscope (Eclipse, Ti) with a 60X oil immersion objective (NA 1.4). During time-lapse imaging, cells plated on 4 well culture chambers (Sarstedt or Ibidi) were kept in an acrylic chamber at 37°C in 5% CO2. Light intensity of each channel was set at 4, with an exposure time of 300 ms and frame interval of 2 sec for 5min. For imaging fixed cells imaging, light intensity of each channel was set at 1, with an exposure time of 100-900 ms for all the fluorophores except for Hoechst dye (5ms). Images were captured with a CoolSNAP HQ2camera (Roper Scientific) using NIS-Elements AR software (version 4.20.01 from Nikon Corporation).

### Long-term live imaging

For long-lasting time-lapse experiments, neurons were stored in an automated incubator/imaging system (Cytation™ 5 Cell Imaging Multi-Mode Reader associated with a BioSpa™ 8 Automated Incubator, BioTek, USA). The plates were stored in the BioSpa at 37°C and 5% CO_2_ and were automatically transferred to the Cytation™ 5 for imaging. For all live-imaging experiments a 20X Plan 0.45 NA objective was used and several fields of view were acquired in time-lapse mode using the point-visiting function.

### STED microscopy and deconvolution

All STED z-stacks were acquired on a *Leica* TCS SP8 gated STED system (*Leica microsystems, Manheim, Germany*) equipped with a pulsed 775 nm depletion laser and a pulsed white light laser (WLL) for excitation. For acquiring images, either a *Leica* Objective HC APO CS2 100×/1.40 Oil or a Glycerol objective (*Leica,* HC APO 93x/1.30 GLYC motCORR) were used.

Rat hippocampal neurons co-labelled with Primary antibodies: Mouse acetylated Tubulin (1:500), Rat Tyrosinated Tubulin (1:500), Rabbit Pericentrin (1:500) followed by species specific secondary antibodies: anti-mouse Abberior Star 580 (1:200; Abberior GmbH Gottingen, Germany), anti-rat Abberior Star RED (1:200; Abberior GmbH Gottingen, Germany), anti-rabbit AF 488 (1:500, Invitrogen) respectively were embedded in Mowiol or Aberrior liquid Mount and excited via the WLL at 640, 561 and 488nm, respectively. Emission was acquired between 650-710nm for Abberior Star RED and 580–620nm for Abberior Star 580 and 500–530nm for AF 488. STED channels. Abberior Star RED and Star 580 were depleted with 50% and 100% with 775 nm depletion laser, respectively. The detector time gates for both channels were set to 0.5– 6ns. The imaging format for all images was set to 2048 × 2048 and an optical zoom of 3 resulted in a pixel size for oil: x/y 18,9nm, for glycerol: x/y 20,4nm. Z spacing was set to 120nm or 160nm. Scan speed was set to 600 lines per second and 8-times line averaging was applied. Confocal overview images were acquired with the same objective, same optical settings but less zoom, less averaging and less excitation power.

Deconvolution of STED z-stacks were done with Huygens Professional (*Scientific Volume Imaging, Hilversum, The Netherlands*). Within the *Deconvolution wizard*, images were subjected to automatic background correction (*lowest* value method), *Signal-to-noise ratio* was set to 15 for both channels and the *Optimized iteration mode of the CMLE* was applied until the algorithm reached 25 iteration steps.

### Confocal Spinning disk imaging

Images were taken with 10X and 60X oil (NA 1.4) objectives on a Nikon EclipseTi2 inverted spinning disk microscope equipped with an LED light source (Lumencor^®^ from AHF analysentechnik AG, Germany), a spinning disk confocal unit (X-Light V2 L-FOV from CrestOptics S.p.A. Italy) and a digital CMOS camera (ORCA-Flash4.0 V3 C13440-20CU from Hamamatsu) controlled with NIS-Elements software. 60X Z-series images with a step size of 300 nm and 1 µm for primary neurons and embryonic brain slices, respectively.

### EB3 comets tacking in the soma of developing neurons

TrackMate (v6.0.1), an open-source Fiji (imageJ) plugin ^47^ was used for the semi-automated tracking EB3 comets from 2D epifluorescence time-lapse live-images of EB3-tDTomato expressing primary neurons. EB3-tdTomato transfected DIV1-2 rat hippocampal neurons (via Amaxa nucleofection) and DIV1 mouse cortical neurons co-transfected with EB3-tdTomato or control or Cep120 shRNA or Cep120 GFP plasmids (via IUE) were used for analysis.

Pre-processing of the EB3 time-lapse images before loading them on to TrackMate: Substract background tool in the Fiji was chosen, Rolling ball radius selection of 1 pixel delineated the EB3 comets from the background distinctively. In order to study the EB3 dynamics in the soma of the developing neurons, we have specifically chosen the neuronal soma area for our analysis. After loading the time-lapse to the TrackMate, LOG detector was selected, and the following parameters were given to be able to detect most but specific EB3 comets in the time-lapse: i). blob diameter was chosen to 3 pixels (∼0.45 microns), ii). threshold value between 25 and 40 (based on the signal to noise ratio). iii). Median filter and sub-pixel localization parameters were set to on. In the next step, no initial thresholding was performed and continued to Hyperstack displayer, this step detects all the comets in the 151 frames (2 frames per sec for 5min). Next, to create tracks from the EB3 comets in the time-lapse, LAP tracker (Jaqaman et al 2008, Nature methods) was selected, under which the following parameters were set without featuring any penalties: a). for Frame to frame linking of the comets track, maximum distance set was to 1 micron; b). for Track segment gap closing, gap closing was allowed and maximum distance set to 1 micron with a maximum frame gap set to 2; c). Track merging was allowed when the maximum distance is 1 micron. The EB3 tracks thus created were checked one-by-one manually using the TrackScheme option. Individual tracks created in the TrackScheme and the actual EB3 comet tracks were verified manually in the time-lapse, false and non-specific tracks were edited. Using the Analysis option all the data related to the EB3 comets and tracks were obtained as .csv files from which the following parameters were analyzed to study the EB3 dynamics (as illustrated in the graph below): 1. EB3 track speed (microns per sec); 2. Growth (displacement) of each EB3 comet per frame (microns) and 3. Total duration of each EB3 tracks (in seconds). To plot the EB3 tracks and comets dynamics near the MTOC, the XY coordinates of the MTOC for each cell is obtained through ImageJ/Fiji by identifying the XY coordinates of the EB3 asters in the time-lapses and set them to (0,0) and then the coordinates of the EB3 tracks’ and comets’ XY coordinates (available in the data obtained after TrackMate analysis) were normalized accordingly.

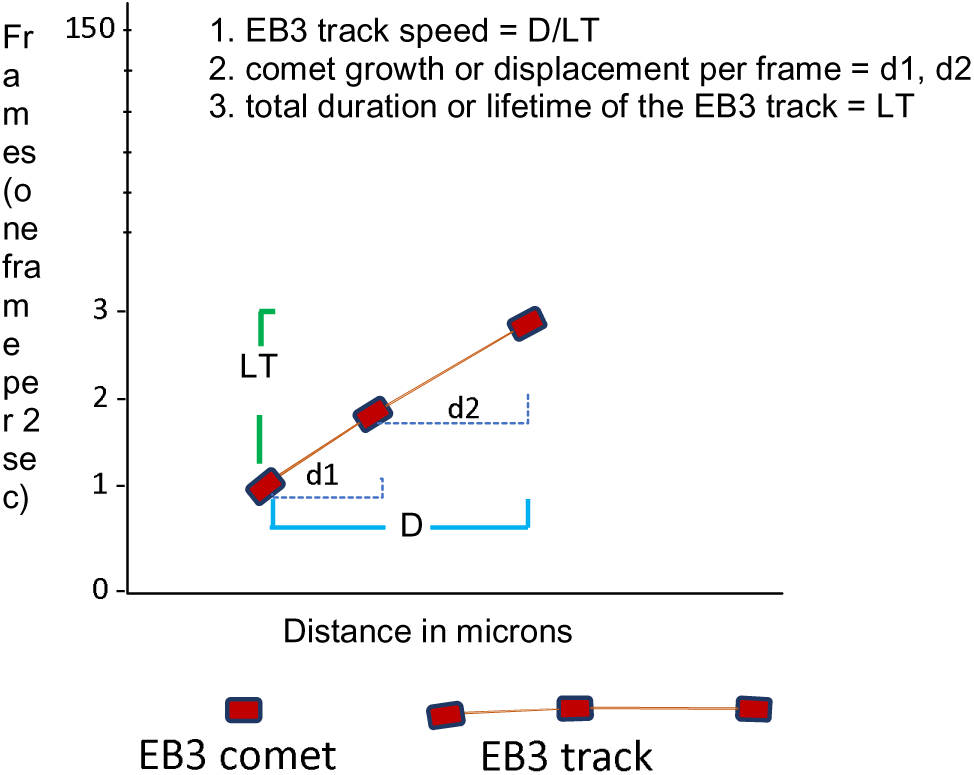

### Axonal phenotype analysis

Epifluorescence 60X oil objective images of Tau-1 or SMI 31 or Ankyrin-G immunostained primary mouse cortical and rat hippocampal neurons were used. Images were loaded onto ImageJ and Tau1/SMI 31 positive gradients in 48-72 hours old neurons and Ankyrin-G rich axon initial segment (AIS) in 7-8 days old neurons, hallmarks of axonal identities, were manually quantified.

### Fluorescent intensity measurements

Images acquired on STED (Tyrosinated tubulin and Acetylated tubulin immunostainings) or confocal spinning disk (Acetylated tubulin immunostainings) microscope were used for analysis. Images were loaded onto ImageJ and z-projection (sum slices) for the entire cell in z-axis was performed on the STED/confocal images. Using ImageJ, soma region was carefully delineated and integrated density in the soma (soma area * mean intensity) was measured. For background correction, mean intensity (background mean intensity) was obtained from the neighboring region (out of the cell). Using the following equation, we obtained the corrected values.

corrected value = integrated density in the soma – (background mean intensity * soma area)

### Neurite length and terminals analysis

Mouse cortical neurons transfected with tDimer together with control, Cep120-shRNA, Cep120-GFP, TACC3-shRNA, SplB-treated control cells and SplB-treated Cep120-GFP cells were used for neurite length and terminals analysis. For Neurite length analysis, in each cell total length of all neurites, length of the longest neurite and length of other neurites were measured manually from all the mentioned conditions using the ROI manger in ImageJ (NIH). For Neurite terminals analysis, total number of neurite terminals per cell was counted manually from all the above-mentioned conditions.

### Cortical cells preparation and FACS sorting

Brain cortices transfected with Cep120 shRNA or controls in combination with pNeuroD-GFP, via IUE at E15, harvested at E19 were prepared as described above. The concentration of shRNA plasmids injected was 4-fold higher than that of the pNeuroD-GFP plasmids. We used 2 µg/µl for shRNA (control or Cep120), 0.5 µg/µl pNeuroD-GFP plasmid. Four days later pregnant mice were anesthetized with CO2/O2, euthanized before taking the E19 embryos out from their uteri. Embryos were then decapitated, skulls were opened, brains were collected in petri dishes with Hibernate^TM^-E medium (Invitrogen) on ice. Hemispheres were separated, meninges were carefully stripped away, and cortices were dissected on ice. Transfected (fluorescent) cortical regions were identified and dissected on ice under a stereo microscope (Olympus SZX16) equipped with a UV light source. The isolated cortical regions were then incubated in 1x HBSS (Invitrogen) containing papain and DNase (Worthington) for 10 min at 37°C neurons, DMEM/10% FCS was added to stop the digestion reaction, then washed with 1x HBSS (warm), which is then replaced with FACS buffer (0,2mM EDTA in Ca-/Mg-free PBS). Cells were then triturated using a fire polished Pasteur pipette with 1mm opening in 2ml FACS buffer for 10-20 times thoroughly but gently until suspended. Cells were then centrifuged at 150g for 10 min in 1 ml cold FACS buffer. Cells were then passed through a 40µm insert filter to remove clusters and filter washed 2 times with 0.5ml FACS buffer. The filtrate was then taken for sorting GFP+ cells at the Cytometry & Cell Sorting Core Unit, Stem Cell Transplant Clinic, Oncology Center at the UKE Hamburg, using the BD FACSAria (TM) Fusion Cell Sorter through a 70 μm Nozzle at 4°C. Immediately after sorting, cells were collected by centrifugation (150-200xg, 10min, 4°C), after which an excess of FACS buffer is removed in sterile conditions. The cells pelleted were then stored in - 80°C until further use. FAC sorted pNeuroD-GFP cells from cortices of 1 or 2 embryos were pooled for proteome analysis.

### Sample preparation for proteome analysis

Ten FACS sorted cell collections were lysed in 100 mM triethyl ammonium bicarbonate (TEAB) and 1% w/v sodium deoxycholate (SDC) buffer, boiled at 95 °C for 5 min and sonicated with a probe sonicator. Disulfide bonds were reduced with dithiothreitol (DTT), alkylated in presence of iodoacetamide (IAA) and digested with trypsin (sequencing grade, Promega) at 37 °C overnight. SDC was precipitated by the addition of 1% v/v formic acid (FA) followed by centrifuged at 16,000 g and the supernatant was transferred into a new tube. Samples were dried in a vacuum centrifuge. Afterwards, samples were resuspended in 100 mM TEAB and labelled with 10-plex TMT according to the manufacture’s instruction, except a 1/10 downscaling was used for labelling low amounts of proteins. After quenching of TMT labelling with hydroxyl amine, the nine samples were combined based on similar cell numbers and dried again in a vacuum centrifuge.

About 1µg each of the TMT labelled samples were resuspended in 10 mM ammonium bicarbonate (pH = 8.5) for basic reversed phase chromatography. Peptides were separated within a 25 min gradient (3 – 35% acetonitrile) on a monolith column (ProSwift™ RP-4H, 1 mm x 250 mm, Thermo Fisher Scientific) using an HPLC system (Agilent 1200 series, Agilent Technologies) with a two-buffer system. Equilibration buffer: 10 mM ammonium bicarbonate (pH = 8.5), elution buffer 10 mM ammonium bicarbonate in 90% acetonitrile. A fraction collector was used to collect 1 min fractions which were then combined to 13 fractions and dried in a vacuum centrifuge.

### Analysis of the TMT-labeled tryptic peptides with liquid chromatography coupled to tandem mass spectrometry (LC-MS/MS)

TMT-labeled tryptic peptides were resuspended in 0.1% formic acid. NanoLC chromatographic separation was achieved on an UPLC system (Dionex Ultimate 3000, Thermo Fisher Scientific)). Attached to the UPLC was a C18 reversed phase peptide (RP) trap (Acclaim PepMap 100, 100 µm x 2 cm, 100 Å pore size, 5 µm particle size) for desalting a purification followed by a C18 RP analytical column (Acclaim PepMap 100, 75 µm x 50 cm, 100 Å pore size, 2 µm particle size). Peptides were separation using a 60 min gradient with increasing ACN concentration from 2% - 30% ACN. The eluting peptides were analyzed on a quadrupole, orbitrap, ion trap tribrid mass spectrometer (Fusion, Thermo Fisher Scientific) in data dependent acquisition (DDA) mode with synchronous precursor selection MS³ (SPS-MS³,^48^). Therefore, the topmost intense ions per precursor scan (2×10^5^ ions, 120,000 Resolution, 120 ms fill time) were analysed first by MS/MS in the ion trap (CID at 35 normalized collision energy, 1×10^4^ ions, 50 ms fill time) in a range of 400 – 1200 m/z and spectra demonstrating TMT reporter ions were further analysed by synchronous precursor selection from the MS/MS spectrum (max. 10) and in the Orbitrap (HCD 65 normalized collision energy, 5×10^4^ ions, 50.000 resolution, 86 msec fill time). A dynamic precursor exclusion of 30 s was used.

### LC-MS/MS Data analysis and processing

Acquired LC-MS/MS data were searched against the mouse (release April 2020, 17,013 protein entries) protein data base downloaded from https:\\www.Uniprot.org (EMBL) using the Sequest algorithm integrated in the Proteome Discoverer software version 2.4 (Thermo Fisher Scientific. Mass tolerances for precursors was set to 10 ppm and 0.6 Da for fragments. Carbamidomethylation on cysteine residues and TMT label modification on the peptide N-terminus and the amine group in the lysine sidechain were set as a fixed modification and the oxidation of methionine, pyro-glutamate formation at glutamine residues at the peptide N-terminus as well as acetylation of the protein N-terminus, methionine loss at the protein N-terminus and the acetylation after methionine loss at the protein N-terminus were allowed as variable modifications. Only peptides with a high confidence (false discovery rate < 1% using a decoy data base approach) were accepted as identified. TMT reporter areas were extracted from corresponding MS³ spectra for each identified peptide and to generate relative protein abundance level across all the 10 samples.

Out of the 10 samples (5x Controls and 5x Cep 120shRNA) used for TMT-labelling, one of the control sample (TMT9) that had low number of FAC sorted GFP+ cells in comparison to the other samples showed high variability for many proteins, hence excluded from the statistical analysis. The least cell number was 12164 GFP+ cells, as estimated by cell sorter, for the rest of the 9 samples (4 x controls and 5 x Cep120 shRNA) used for further analysis.

### Bioinformatic analysis of proteome data

For heatmap, a hierarchical clustering of the 50 proteins with the lowest significant p-value (< 0.05) that identifies a relative down regulation of protein expression in Cep120 shRNA neurons in comparison with the controls from normalized data as stated in the Raw data (See Supplementary excel file Rawdata.xlsx for details). The dendrogram represents the hierarchical clustering with a distance score based on Pearson-correlation coefficients (z-score -2 to 2) and a complete linkage clustering as cluster agglomeration criteria ((dis-)similarity between group Cep120 versus Control).

For the String analysis, the proteins whose levels were found to be significantly different between the two conditions (See Supplementary excel file STRING.xlsx for details)., were used STRING v11 ^49^, allowing to predict functional associations between proteins ^50^.

Gene set enrichment analysis (GSEA) was performed with WebGestalt ^51^. Briefly all the proteins that were quantified in each of the replicates (N=1093) were ordered by their relative difference between the two conditions and analyzed with WebGestalt for the three non-redundant functional gene ontology databases (biological process, cellular component, molecular function). All the advanced parameters were kept as indicated in the presets (for details refer to http://www.webgestalt.org/). (See Supplementary excel file GSEA.xlsx for details).

### Image processing

Linear adjustment of brightness and contrast was performed on images using Photoshop CS or ImageJ.

### Statistical analysis

Statistical analysis was performed using the GraphPad Prism 8 software. The Student’s t-test (two-tailed) was used to compare means of two groups, whereas analysis of variance (ANOVA) was used when comparing more than two groups. Asterisks *, **, *** and **** represent P < 0.05, 0.01, 0.001, and 0.0001, respectively. Error bars in the graphs always represent standard error of mean. Pearson correlation analysis was performed for showing a linear relationship between two sets of data.

## Acknowledgements

We thank M. Richter (Calderon lab, ZMNH, UKE), I. Hermans-Borgmeyer (ZMNH, UKE), and the members of animal facility at the UKE Hamburg for their help with animal experiments. Thanks to the members of Cytometry & Cell Sorting Core Unit, Stem Cell Transplant Clinic, Oncology Center at the UKE Hamburg for their help with FACS experiments.

Funding Sources: FCA is supported by Deutsche Forschungsgemeinschaft (DFG) Grants: FOR 2419, CA1495/1-1, and CA 1495/4-1; ERA-NET Neuron Grants (Bundesministerium für Bildung und Forschung, BMBF, 01EW1410, and 01EW1910), JPND Grant (Bundesministerium für Bildung und Forschung, BMBF, 01ED1806), and University Medical Center Hamburg-Eppendorf (UKE). DPM is a co-applicant in the DFG Grant CA 1495/4-1 for FCA. EFF is supported by a Schram Stiftung (T0287/35359/2020) and a DFG grant (FO 1342/1-3).

## Author contributions

FCA conceived the idea, supervised the project, and wrote the manuscript. FCA and DPM designed research and interpreted the data. BS conducted all the cell culture work. OK and DPM performed STED imaging, OK did the deconvolution and FCA performed the analysis. DPM and FCA performed all *in utero* electroporation surgeries. DPM, NS, SW, BS and RR performed all the immunostainings. DPM and NS did all the epifluorescence fixed cell imaging; DPM, SW, NS and FCA did the analysis. DPM performed epifluorescence time-lapse imaging and TrackMate semi-automated analysis of EB3 comets. RR and DPM performed cryosectioning of embryonic mouse brains. DPM did the spinning disk confocal imaging and analysis for primary neurons and for *in situ* experiments. EFF performed the long-term epifluorescence time-lapse acquisitions, *post hoc* SMI 31 immunostainings and imaging; DPM quantified the data. BS and DPM prepared cells for FAC sorting. HS designed and supervised the LC-MS/MS experiment: CK did the experimental work, analysis and processing. EFF performed the final bioinformatic analysis of proteome data, TR worked on preliminary proteome data analysis and coordination with FACS facility. All authors provided input for writing the manuscript.

## Supplementary Figure legends

**Supp. Figure 1.**
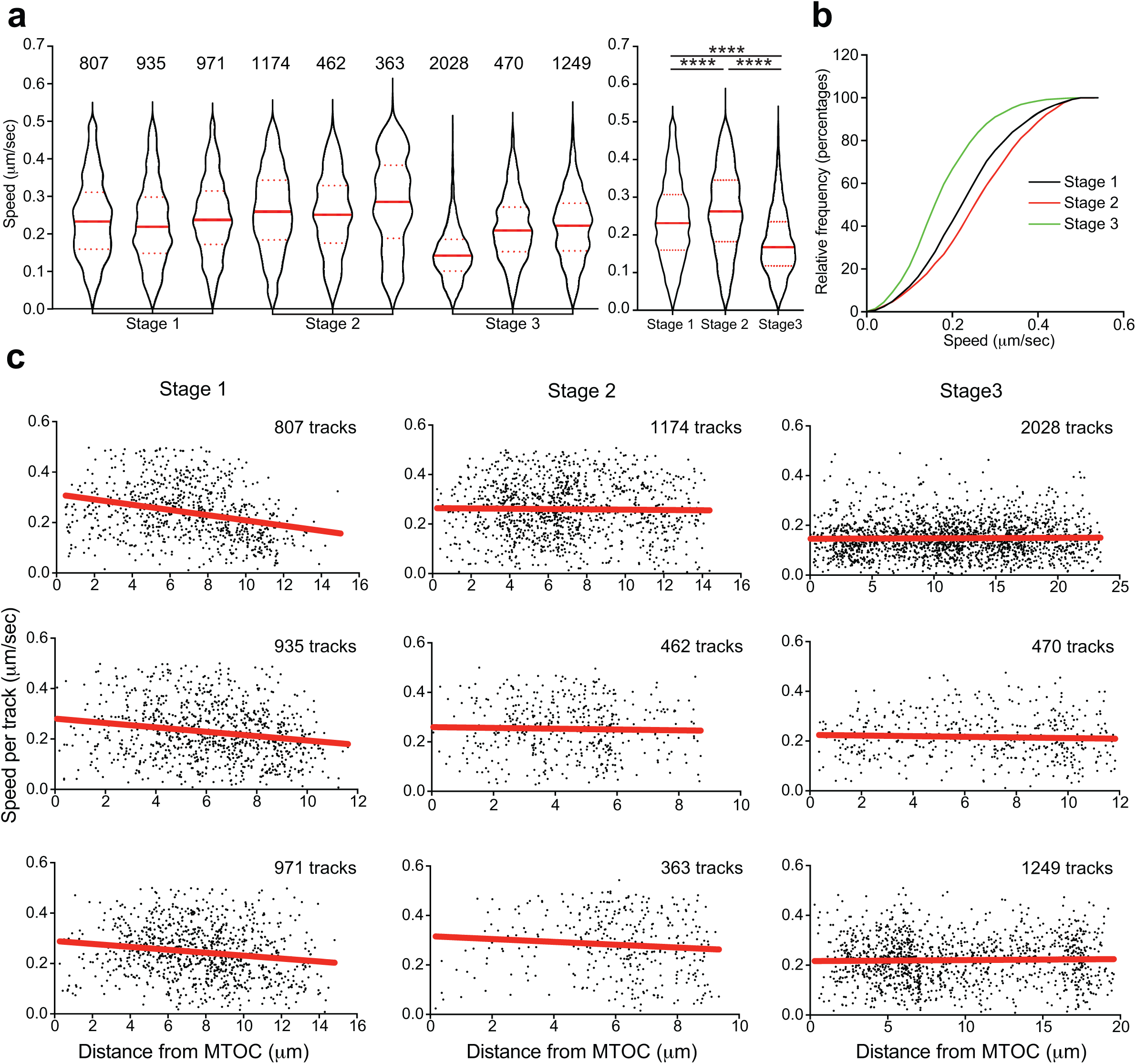
EB3 track growth speed is higher in the soma of multipolar neurons (related to Figure 2) **a.** EB3 track growth speed (µm/sec) in the soma of developing neurons. Violin plots show distribution of EB3 track growth speed values in 3 cells individually (left panel; insets show no. of EB3 tracks analyzed per cell) and combined (Right panel) for stage 1, stage 2 and stage 3 neurons used for analysis in Figure 2 **a-c**. Median (thick red line), 25 and 75% quartiles (dotted red lines) were shown. Mean ± SEM values of EB3 growth speed combined from all stage 1 cells = 0.2373 ± 0.0020, stage 2 = 0.2622 ± 0.0025, stage 3 = 0.1807 ± 0.0014. P < 0.0001 by one-way ANOVA, post hoc Tukey’s test, **** P < 0.0001; values from n = 3 cells per stage. **b.** Cumulative frequency distribution (percentages) of growth speed (µm/sec) of all EB3 tracks in the soma of stage 1, stage 2 and stage 3 neurons used for analysis in panel **a** and in Figure 2 **a-c**. **c.** Linear regression histograms of growth speed (µm/sec) all EB3 tracks plotted against the distance from the MTOC are plotted for 3 cells each of stage 1, stage 2 and stage 3 cells. Insets show the number of EB3 tracks analyzed per cell.

**Supp. Figure 2.**
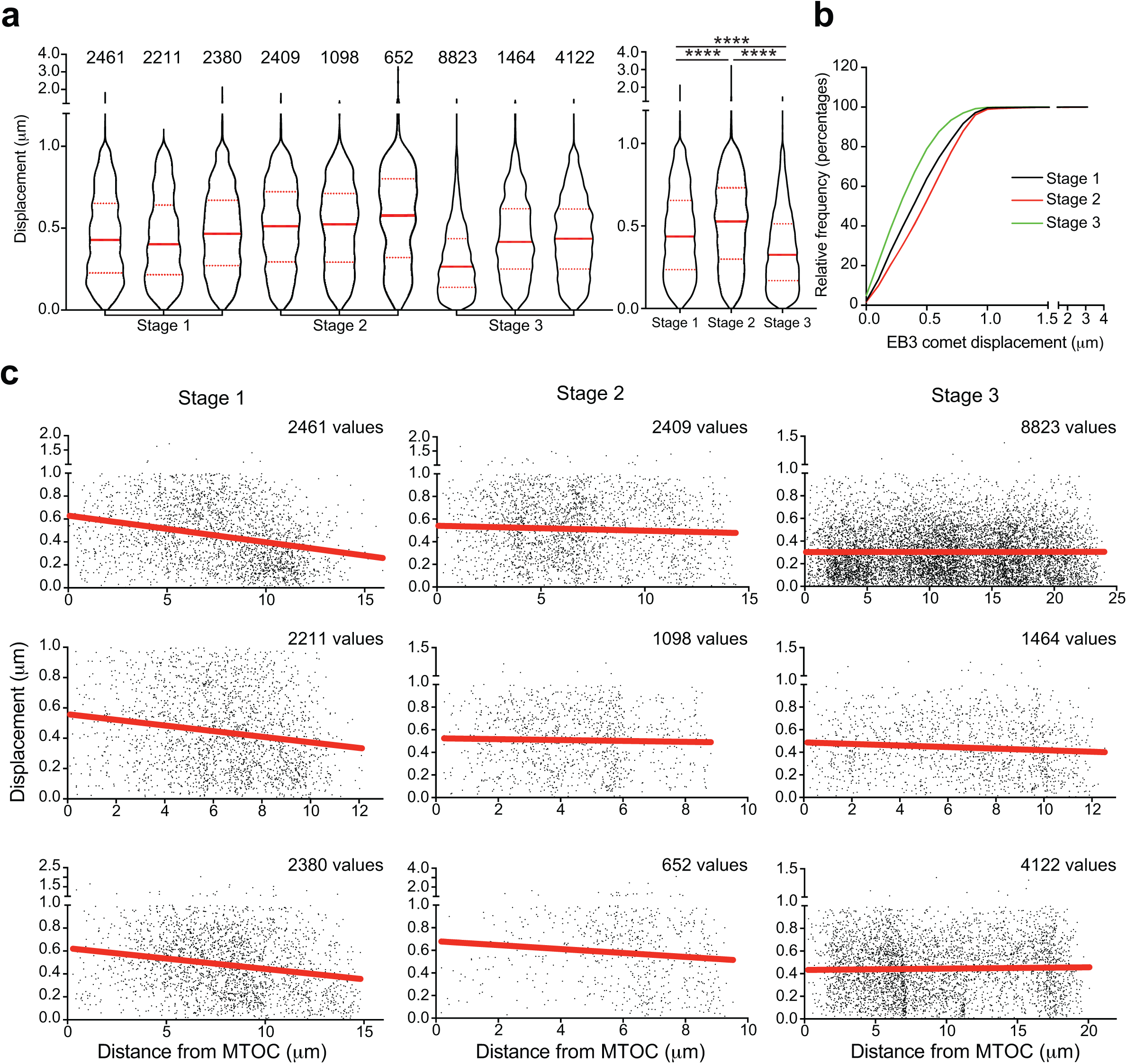
EB3 comet displacement per frame (every 2 sec) is higher in the soma of multipolar neurons (related to Figure 2) **a.** EB3 comet displacement per frame (µm) in the soma of developing neurons. Violin plots show distribution of EB3 comet displacement values in 3 cells individually (left panel; insets show no. of EB3 comets displacement values analyzed per cell) and combined (Right panel) for stage 1, stage 2 and stage 3 neurons used for analysis in Figure 2 **a-c**. Median (thick red line), 25 and 75% quartiles (dotted red lines) were shown. Mean ± SEM values of EB3 comet displacement per frame combined from all stage 1 cells = 0.4566 ± 0.0031, stage 2 cells = 0.5206 ± 0.0042, stage 3 cells = 0.3580 ± 0.0019. P < 0.0001 by one-way ANOVA, post hoc Tukey’s test, **** P < 0.0001; values from n = 3 cells per stage. **b.** Cumulative frequency distribution (percentages) of all EB3 comets displacement (µm) per frame in the soma of stage 1, stage 2 and stage 3 neurons used for analysis in panel **a** and in Figure 2 **a-c**. **c.** Linear regression histograms of all EB3 comets displacement per frame (µm) plotted against the distance from the MTOC are plotted for 3 cells each of stage 1, stage 2 and stage 3 cells. Insets show the number of EB3 comets displacement values analyzed per cell.

**Supp. Figure 3.**
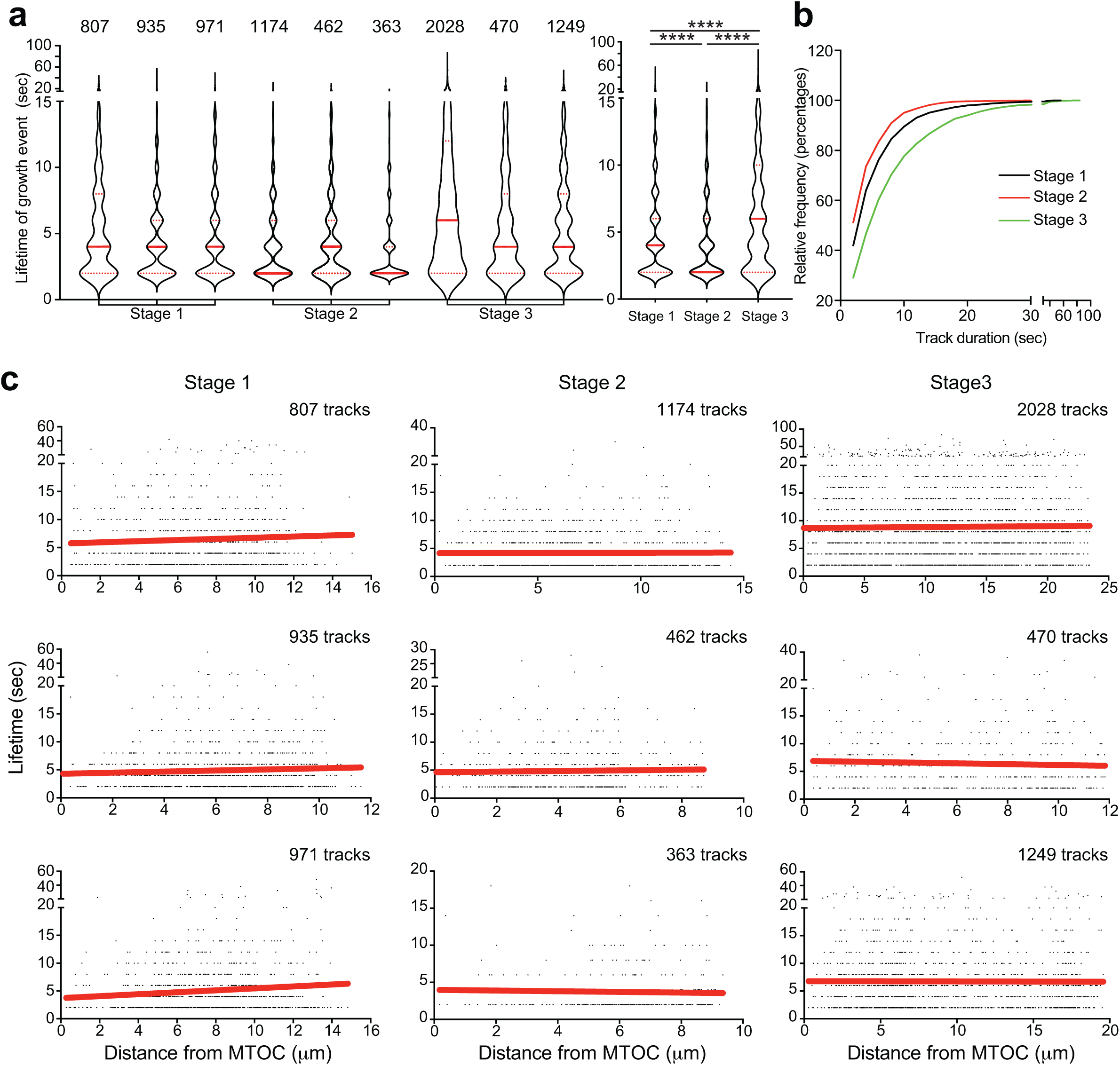
EB3 track duration is higher in the soma of polarized, stage 3, neurons. (related to Figure 2) **a.** EB3 track duration (sec) in the soma of developing neurons. Violin plots show distribution of EB3 track duration values in 3 cells individually (left panel; insets show no. of EB3 tracks analyzed per cell) and combined (Right panel) for stage 1, stage 2 and stage 3 neurons used for analysis in Figure 2 **a-c**. Median (thick red line), 25 and 75% quartiles (dotted red lines) were shown. Mean ± SEM values of EB3 track duration stage 1 cell pooled = 5.415 ± 0.0984, stage 2 = 4.269 ± 0.0768, stage 3 = 7.860 ± 0.1242. P < 0.0001 by one-way ANOVA, post hoc Tukey’s test, **** P < 0.0001; values from n = 3 cells per stage. **b.** Cumulative frequency distribution (percentages) of duration (sec) of EB3 tracks in the soma of stage 1, stage 2 and stage 3 neurons used for analysis in panel **a** and in Figure 2 **a-c**. **c.** Linear regression histograms of all EB3 tracks duration (sec) plotted against the distance from the MTOC are plotted for 3 cells each of stage 1, stage 2 and stage 3 cells. Insets show the number of EB3 tracks analyzed per cell.

**Supp. Figure 4.**
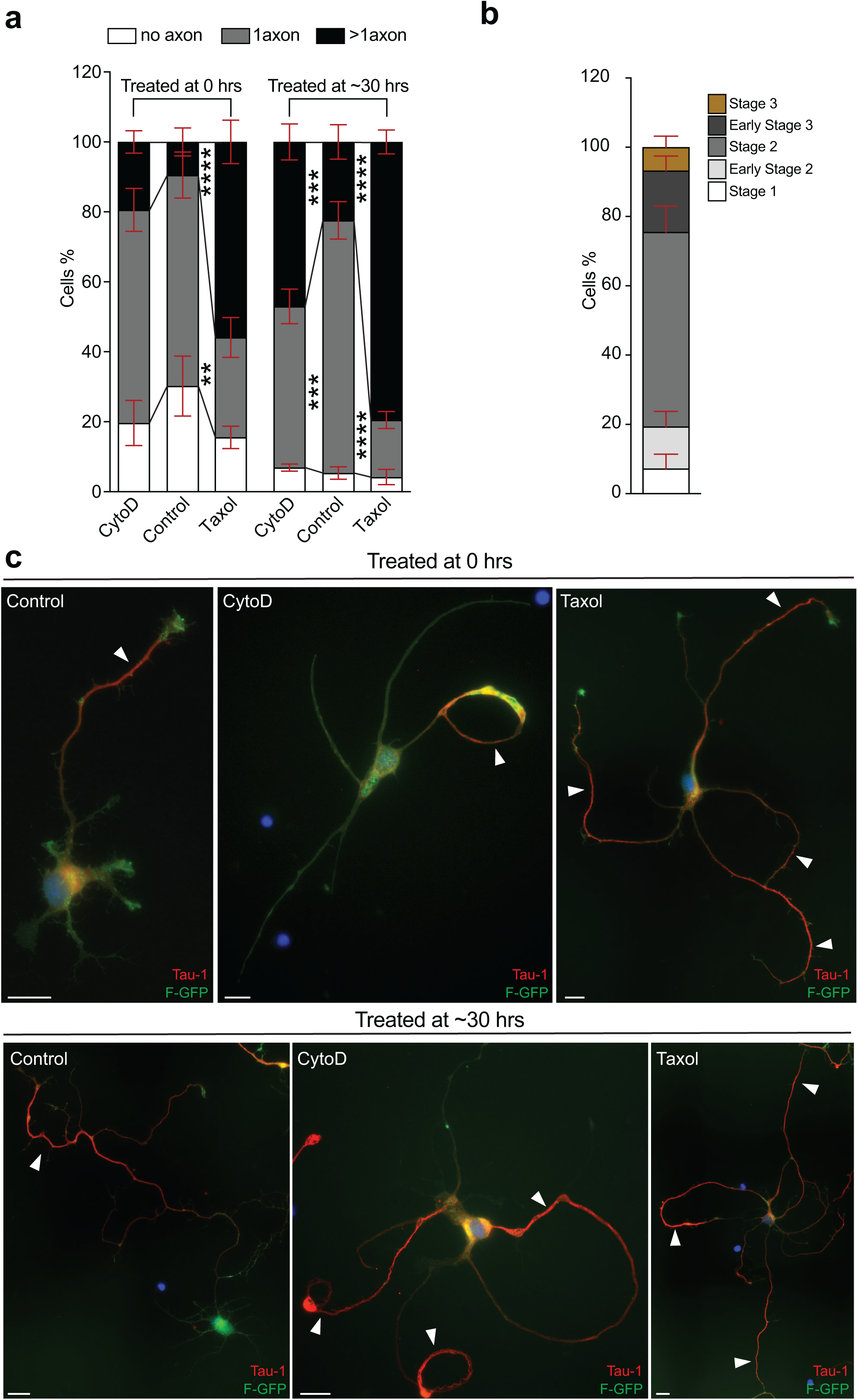
Contrary to microtubule stabilization (by Taxol), actin disruption (by cytochalasin D) induced multipolarity is time-dependent – confirmed by Tau-1 immunostaining. (related to Figure 3) **a.** Quantifications show the percentage of neurons that were either treated immediately (0 hrs.) or ∼ 30 hrs. of plating in vitro with DMSO or 2µM cytochalasin D or 5nM Taxol for 48 hours differentiated to have no axon or 1 axon or more than 1 axon. Mean ± SEM values for percentage of cells treated at 0 hrs. with DMSO: no axons = 30.22 ± 8.554, 1 axon = 60.32 ± 6.606, more than 1 axon = 9.463 ± 3.979; CytoD: no axons = 19.65 ± 6.441, 1 axon = 60.94 ± 6.129, more than 1 axon = 19.42 ± 3.226; Taxol: no axons = 15.56 ± 3.195, 1 axon = 28.54 ± 5.698, more than 1 axon = 55.91 ± 6.196; treated at ∼ 30 hrs. with DMSO: no axons = 5.333 ± 1.781, 1 axon = 72.23 ± 5.354, more than 1 axon = 22.44 ± 4.914; CytoD: no axons = 6.898 ± 1.021, 1 axon = 46.09 ± 4.951, more than 1 axon = 47.02 ± 5.115; Taxol: no axons = 4.180 ± 2.181, 1 axon = 16.34 ± 2.419, more than 1 axon = 79.48 ± 3.403. α = 0.05 by two-way ANOVA, post hoc Tukey’s test, **** P < 0.0001, *** P < 0.001, ** P < 0.01. Data is obtained from 4 different hippocampal cultures. **b.** Quantifications show the percentage of neurons (untreated) after ∼ 30hrs. of plating in vitro in different stages of development, stage 1 to stage 3. **c.** Images of Farnesylated-GFP (F-GFP) transfected neurons that were either treated immediately (0 hrs.) or ∼ 30 hrs. of plating in vitro with DMSO or 2 μM Cytochalasin D or 5nM Taxol for 48 hr. The cells are then PFA fixed for post hoc Tau-1 (shown in red) immunostainings to confirm axonal (indicated by white arrowheads) identity of the neurites. Scale bar: 10 µm.

**Supp. Figure 5.**
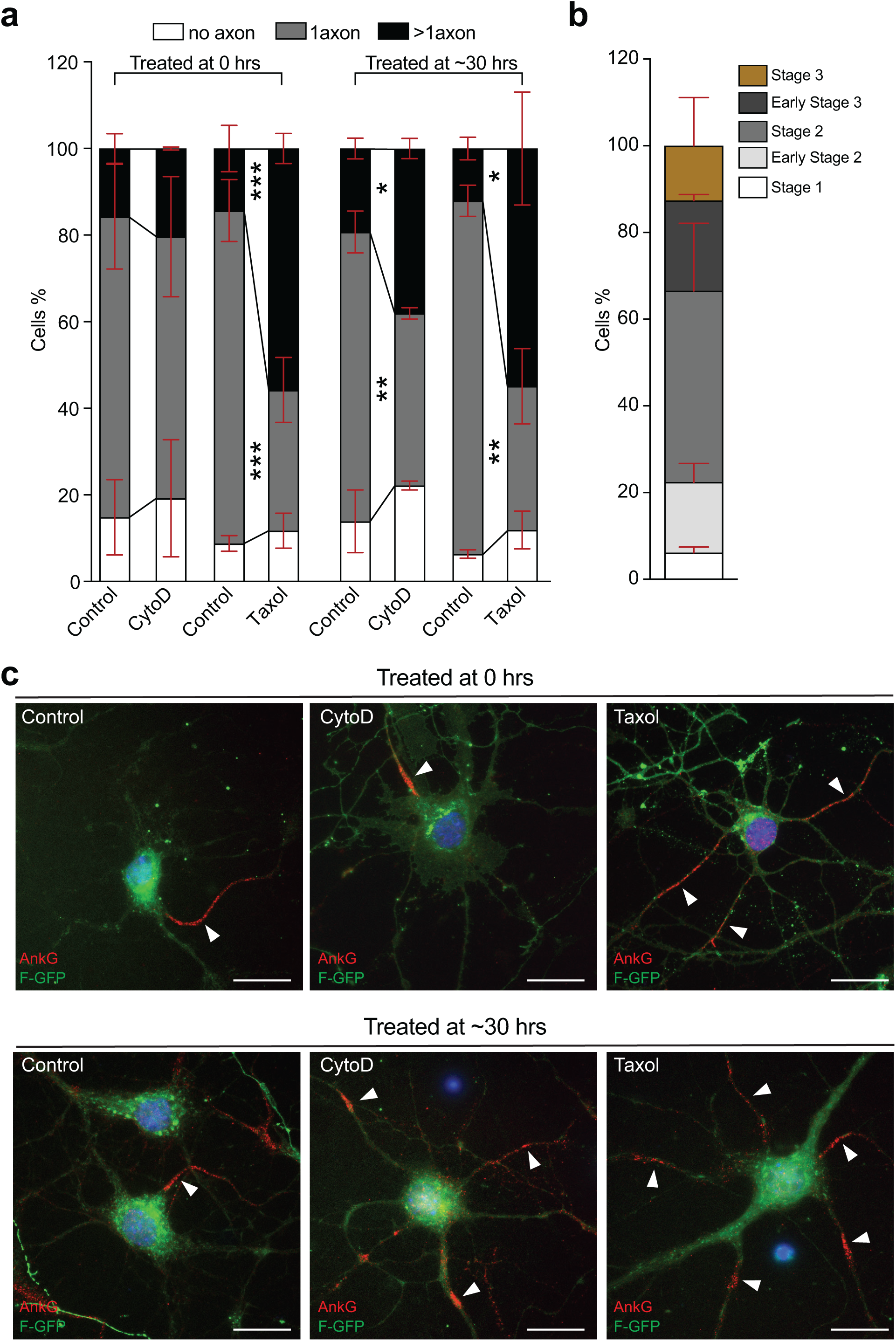
Contrary to microtubule stabilization (by Taxol), actin disruption (by cytochalasin D) induced multipolarity is time-dependent – confirmed by Ankyrin-G immunostaining. (related to Figure 3) **a.** Quantifications show the percentage of neurons that were either treated immediately (0 hrs.) or ∼ 30 hrs. of plating in vitro with DMSO or 2µM cytochalasin D or 5nM Taxol for 7 days differentiated to have no axon or 1 axon or more than 1 axon. For cytochalasin D treatment group cells, fresh medium was exchanged after 16 hours to prevent cells from dying otherwise when incubated with cytochalasin D for 7days. Mean ± SEM values for percentage of cells treated at 0 hrs. with DMSO (CytoD controls): no axons = 14.84 ± 8.690, 1 axon = 69.45 ± 12.10, more than 1 axon = 15.72 ± 3.405; CytoD: no axons = 19.24 ± 13.53, 1 axon = 60.42 ± 13.87, more than 1 axon = 20.35 ± 0.345; with DMSO (Taxol controls): no axons = 8.805 ± 1.805, 1 axon = 76.85 ± 7.150, more than 1 axon = 14.35 ± 5.350; Taxol: no axons = 11.74 ± 4.050, 1 axon = 32.50 ± 7.500, more than 1 axon = 55.76 ± 3.450. Mean ± SEM values for percentage of cells treated at ∼ 30 hrs. Mean ± SEM values for percentage of cells treated at 0 hrs. with DMSO (CytoD controls): no axons = 13.90 ± 7.230, 1 axon = 66.82 ± 4.850, more than 1 axon = 19.29 ± 2.385; CytoD: no axons = 22.18 ± 1.030, 1 axon = 39.77 ± 1.305, more than 1 axon = 38.05 ± 2.335; with DMSO (Taxol controls): no axons = 6.380 ± 0.9700, 1 axon = 81.54 ± 3.600, more than 1 axon = 12.09 ± 2.625; Taxol: no axons = 11.92 ± 4.365, 1 axon = 33.20 ± 8.665, more than 1 axon = 54.89 ± 13.03. α = 0.05 by two-way ANOVA, post hoc Sidak’s test, *** P < 0.001, ** P < 0.01, ** P < 0.05. Data is obtained from 2 different hippocampal cultures. **b.** Quantifications show the percentage of neurons (untreated) after ∼ 30hrs. of plating in vitro in different stages of development, stage 1 to stage 3. **c.** Images of Farnesylated-GFP (F-GFP) transfected neurons that were either treated immediately (0 hrs.) or ∼ 30 hrs. of plating in vitro with DMSO or 2 μM Cytochalasin D or 5nM Taxol for 48 hr. The cells are then PFA fixed for post hoc Ankyrin-G (shown in red) immunostainings to confirm axonal (indicated by white arrowheads) identity of the neurites. Scale bar: 10 µm.

**Supp. Figure 6.**
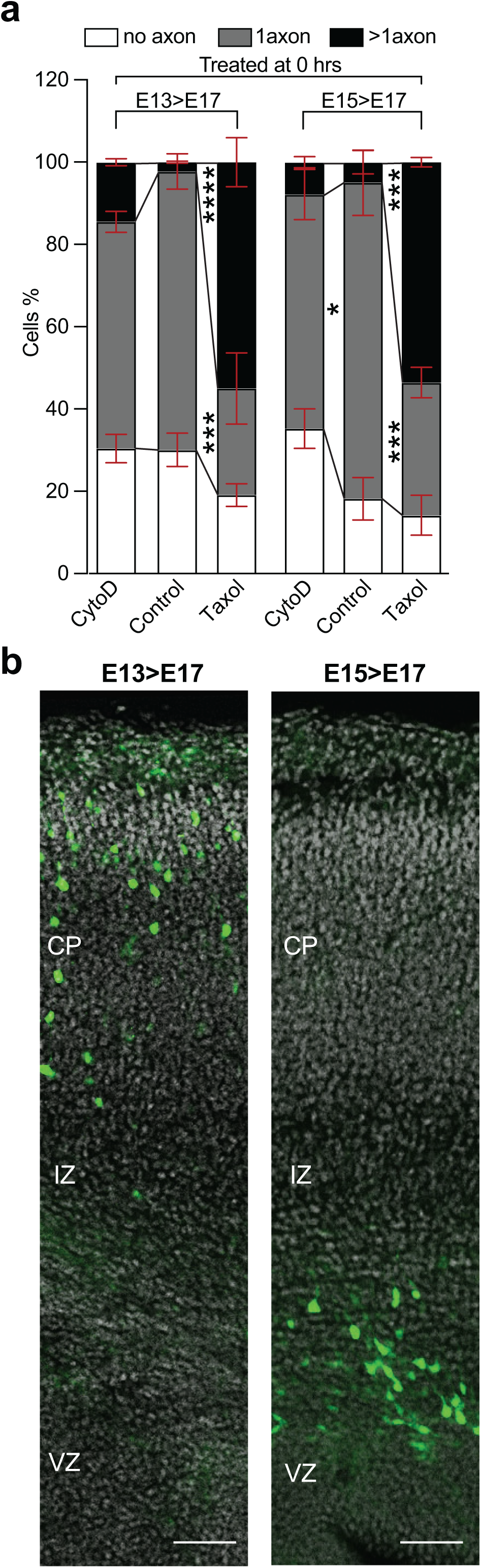
Time-dependent effect of actin disruption (by cytochalasin D) is independent of the differentiated state of the neurons at the time of plating: cells in the intermediate zone versus cells in the cortical plate. (related to Figure 3) **a.** Quantifications show the percentage of cortical neurons from E13 versus E15 IUE brains transfected with Venus plasmid cultured at E17 that were treated immediately (0 hrs.) after plating with DMSO or 2µM cytochalasin D or 5nM Taxol for 7 days differentiated to have no axon or 1 axon or more than 1 axon. Axonal identities in the PFA-fixed cells are confirmed by Tau-1 immunostainings. Mean ± SEM values for percentage of E13 IUE cells treated with DMSO: no axons = 30.08 ± 4.075, 1 axon = 67.71 ± 4.295, more than 1 axon = 2.220 ± 0.2200; CytoD: no axons = 30.41 ± 3.490, 1 axon = 55.10 ± 2.595, more than 1 axon = 14.49 ± 0.8900; Taxol: no axons = 19.13 ± 2.770, 1 axon = 25.88 ± 8.675, more than 1 axon = 55.00 ± 5.905; E15 IUE cells treated with DMSO: no axons = 18.22 ± 5.180, 1 axon = 76.79 ± 7.990, more than 1 axon = 4.985 ± 2.815; CytoD: no axons = 35.22 ± 4.785, 1 axon = 56.92 ± 6.120, more than 1 axon = 7.860 ± 1.340; Taxol: no axons = 14.18 ± 4.875, 1 axon = 32.29 ± 3.715, more than 1 axon = 53.54 ± 1.160. α = 0.05 by two-way ANOVA, post hoc Tukey’s test, **** P < 0.0001, *** P < 0.001, * P < 0.05. Data is obtained from cortical cultures of IUE mouse embryos from 2 different mothers. **b.** E17 mouse brain cortices that were either electroporated at E13 or E15 via IUE with Venus plasmids shows that the E13>E17 brains has more differentiated neurons mostly in cortical plate (CP), the E15>E17 brains has less developed neurons residing in the lower intermediate zone (IZ). Scale bar: 100 µm.

**Supp. Figure 7.**
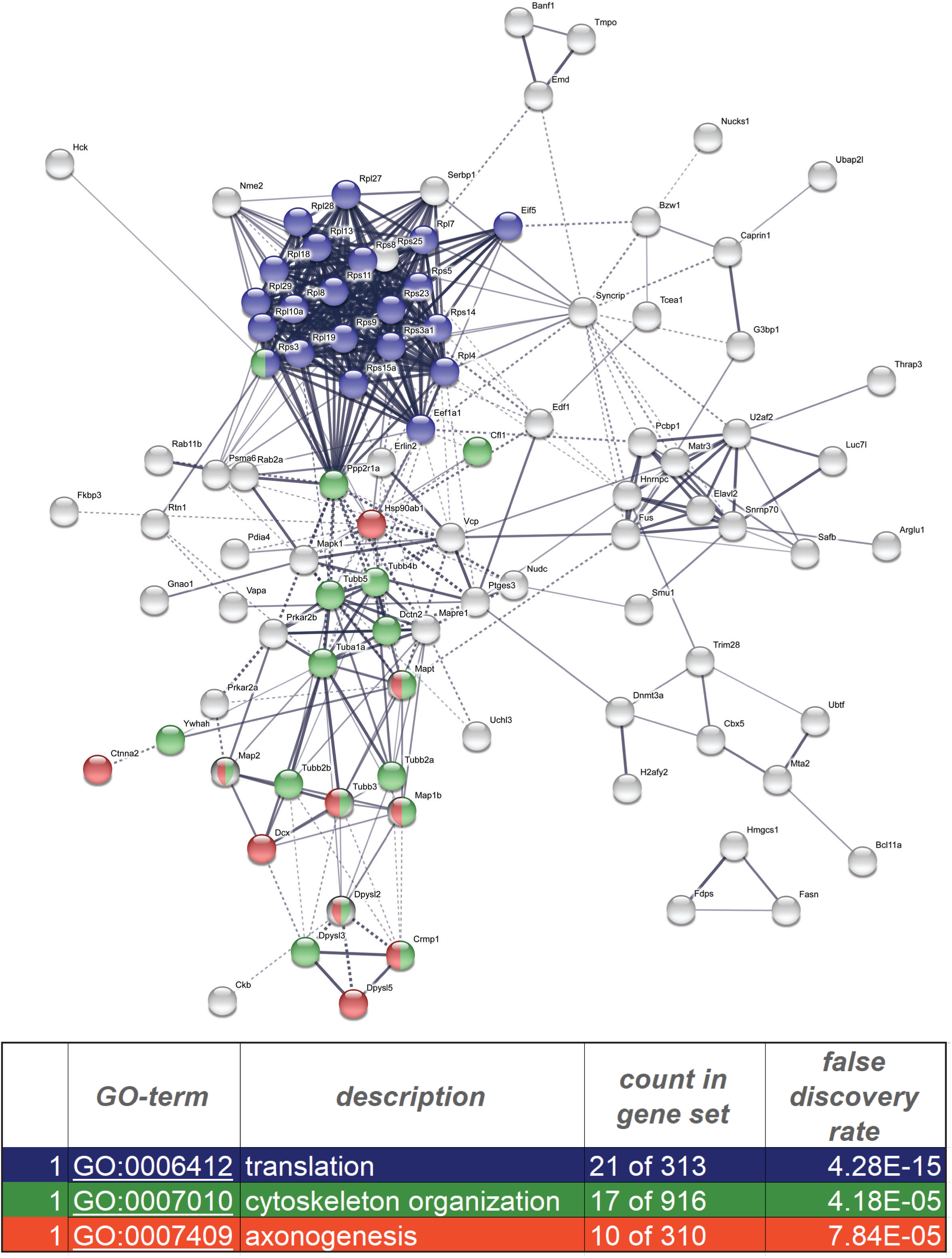
STRING analysis visualization of the proteins that were significantly altered between the control and Cep120 shRNA conditions (P < 0.05). (related to Figure 5) STRING analysis visualization of all the proteins whose levels were significantly changed between the control and Cep120 shRNA conditions (P < 0.05). For simplicity, the disconnected proteins are hidden in the visualization and the “confidence” link visualization is selected (highlighting the strength of data support in each network edge). For further details refer to the original work ^49^. The three following GO terms are colored and detailed as shown in the legend at the bottom of the figure: GO:0007409-axonogenesis; GO:0007010-cytoskeletal organization and GO:0006412-translation. (See Supplementary excel file STRING.xlsx for details).

**Supp. Figure 8.**
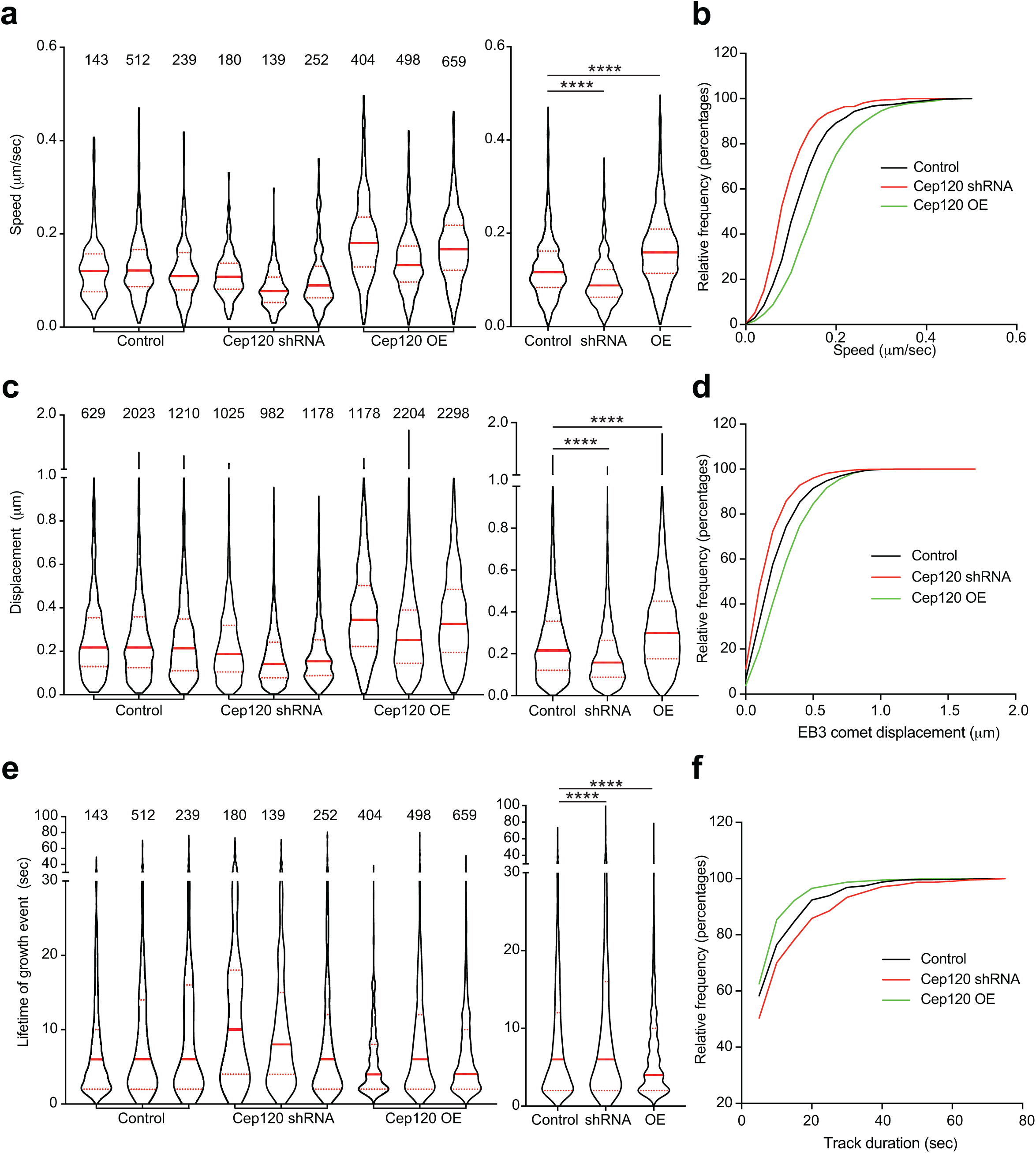
Cep 120 knockdown and overexpression bidirectionally regulates EB3 dynamics in stage 2 neurons. (related to Figure 6) **a.** EB3 track growth speed (µm/sec) in the soma of control, Cep120 shRNA and Cep 120-GFP neurons (shown in Figure 6). Violin plots show distribution of EB3 track growth speed in 3 cells individually (left panel; insets show no. of EB3 tracks analyzed per cell) and combined (Right panel) for control, Cep120 shRNA and Cep120-GFP neurons used for analysis in Figure 6 **a-b**. Median (thick red line), 25 and 75% quartiles (dotted red lines) were shown. Mean ± SEM values of all EB3 track growth speed from the soma of control cells = 0.1294 ± 0.0023, Cep120 shRNA cells = 0.1073 ± 0.0023, Cep120-GFP cells = 0.1677 ± 0.0020. P < 0.0001 by one-way ANOVA; **** P < 0.0001 by post hoc Tukey’s test. Values from n = 3 cells per stage. **b.** Cumulative frequency distribution (percentages) of EB3 track growth speed (µm/sec) in the soma of stage 2 neurons, all values from 3 cells each of control, Cep120 shRNA and Cep 120-GFP groups used for analysis in panel **a** and in Figure 6 **a-b**. **c.** EB3 comet displacement per frame (µm) in the soma of control, Cep120 shRNA and Cep 120-GFP neurons (shown in Figure 6). Violin plots show distribution of EB3 comet displacement per frame values in 3 cells individually (left panel; insets show no. of EB3 comets displacement values analyzed per cell) and combined (Right panel) for control, Cep120 shRNA and Cep120-GFP neurons used for analysis in Figure 6 **a-b**. Median (thick red line), 25 and 75% quartiles (dotted red lines) were shown. Mean ± SEM values of EB3 comet displacement per frame (µm) from the soma of control cells = 0.2626 ± 0.0030, Cep120 shRNA cells = 0.2013 ± 0.0027, Cep120-GFP cells = 0.3330 ± 0.0027. P < 0.0001 by one-way ANOVA; **** P < 0.0001 by post hoc Tukey’s test. Values from n = 3 cells per stage. **d.** Cumulative frequency distribution (percentages) of EB3 comet displacement per frame (µm) in the soma of stage 2 neurons, all values from 3 cells each of control, Cep120 shRNA and Cep 120-GFP groups used for analysis in panel **a** and in Figure 6 **a-b**. **e.** EB3 track duration (sec) in the soma of control, Cep120 shRNA and Cep 120-GFP neurons (shown in Figure 6). Violin plots show distribution of EB3 track duration values in 3 cells individually (left panel; insets show no. of EB3 tracks analyzed per cell) and combined (Right panel) for control, Cep120 shRNA and Cep120-GFP neurons used for analysis in Figure 6 **a-b**. Median (thick red line), 25 and 75% quartiles (dotted red lines) were shown. Mean ± SEM values of all EB3 tracks duration (sec) from the soma of control cells = 9.036 ± 0.3155, Cep120 shRNA cells = 11.33 ± 0.4784, Cep120-GFP cells = 7.358 ± 0.1811. P < 0.0001 by one-way ANOVA; **** P < 0.0001 by post hoc Tukey’s test. Values from n = 3 cells per stage. **f.** Cumulative frequency distribution (percentages) of EB3 track duration (sec) in the soma of stage 2 neurons, all values from 3 cells each of control, Cep120 shRNA and Cep 120-GFP groups used for analysis in panel **a** and in Figure 6 **a-b**.

**Supp. Figure 9.**
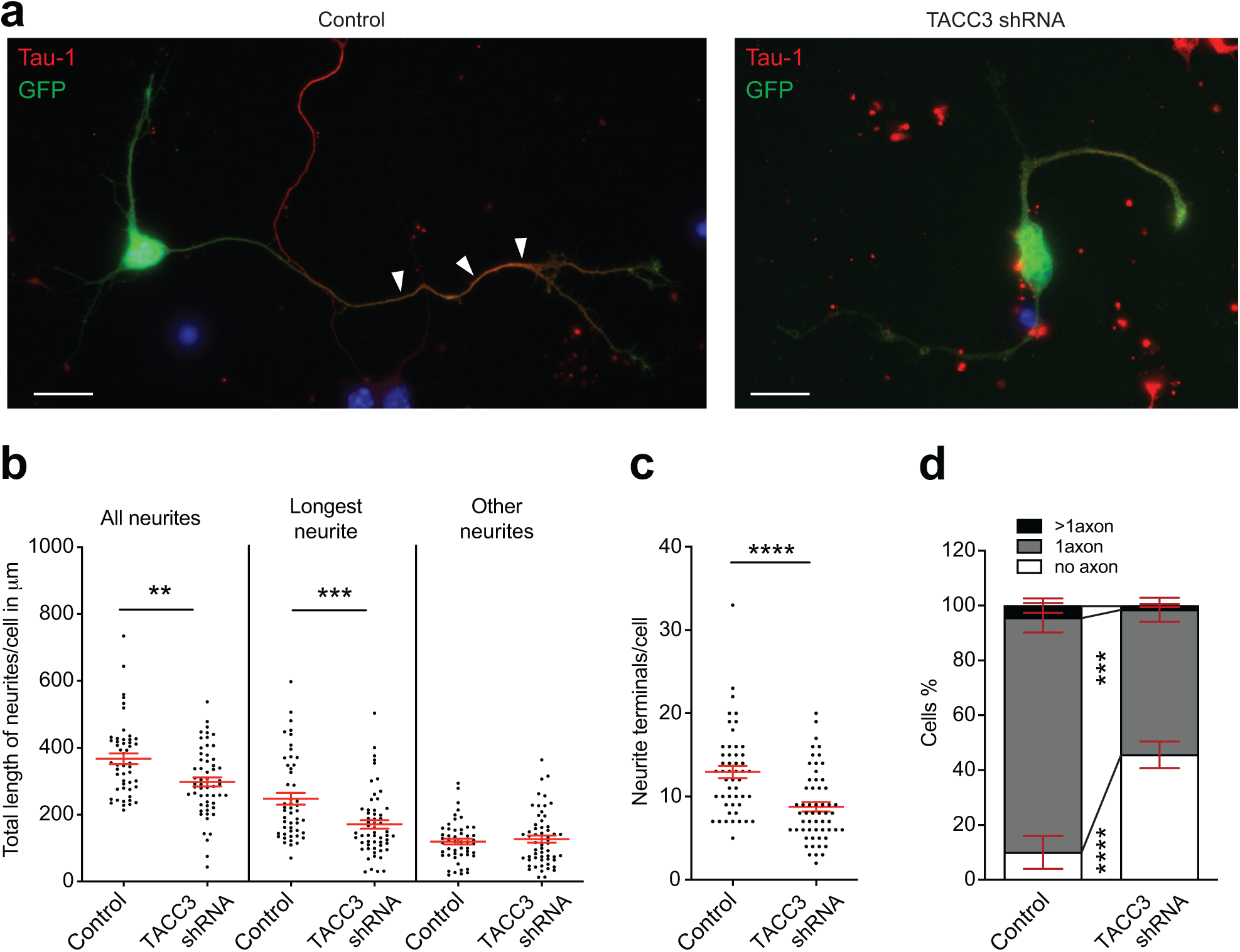
TACC3 downregulation affects axon formation. (related to Figure 7) **a.** Epifluorescence images of mouse cortical neurons transfected via IUE at E15 with control shRNA-GFP or TACC3 shRNA-GFP cultured at E17 for 48 or 72 hrs. immunostained with Tau-1 antibody to confirm axonal identity (indicated by white arrow heads). Scale bar: 10 µm. **b.** Neurite length quantification in neurons (in µm) control shRNA-GFP (n = 50) and TACC3 shRNA-GFP (n = 57) expressing neurons as shown in **a**. Mean ± SEM values for length of all neurites in control cells = 367.1 ± 15.77, TACC3 shRNA-GFP cells = 298.1 ± 13.35. Mean ± SEM values for length of longest neurite in control cells = 247.6 ± 17.60, TACC3 shRNA-GFP cells = 171.0 ± 12.88. Mean ± SEM values for length of other neurites in control cells = 119.4 ± 8.770, TACC3 shRNA-GFP cells = 127.0 ± 10.65. ** P = 0.0011, *** P = 0.0005 by unpaired Student’s t-test. **c.** Neurite terminals quantification in control shRNA-GFP (n = 50) and TACC3 shRNA-GFP (n = 57) expressing neurons as shown in **a**. Mean ± SEM values for neurite terminal per cell in control neurons = 12.94 ± 0.7315, TACC3 shRNA-GFP cells = 8.772 ± 0.5636. **** P < 0.0001 by unpaired Student’s t-test. **d.** Quantifications show the percentage of neurons expressing control shRNA-GFP (n = 225 cells) and TACC3 shRNA-GFP (n = 243 cells), as shown in **a**, differentiated to have no axon, 1 axon and more than 1 axon. Mean ± SEM values for percentage of control cells with no axon = 10.08 ± 5.975, 1 axon = 85.50 ± 5.366, more than 1 axon = 4.430 ± 2.623; percentage of TACC3 shRNA cells with no axon = 45.59 ± 4.835, 1 axon = 52.88 ± 4.406, more than 1 axon = 1.533 ± 0.5436. α = 0.05 by two-way ANOVA, post hoc Sidak’s test, **** P < 0.0001, *** P < 0.001. Data shown in **b**, **c** and **d** is obtained from cortical cultures of IUE mouse embryos from 2 different mothers.

**Video 1.** 3D view of a z-series STED projection of stage 1 rat hippocampal neuron immunostained for tyrosinated tubulin (left) and acetylated tubulin (right).

**Video 2.** Unprocessed (left) and background subtracted (right) 5 minute time-lapse (1 frame every 2 sec) of a EB3-tdTomato-trasfected stage 1 rat hippocampal neuron

**Video 3.** Unprocessed (left) and background subtracted (right) 5 minute time-lapse (1 frame every 2 sec) of a EB3-tdTomato-trasfected stage 2 rat hippocampal neuron,

**Video 4.** Unprocessed (left) and background subtracted (right) 5 minute time-lapse (1 frame every 2 sec) of a EB3-tdTomato-trasfected stage 3 rat hippocampal neuron,

**Video 5.** Long-term time-lapse of a stage 1 rat hippocampal neuron treated with DMSO, PFA fixed after 52 hrs., immunostained for βIII tubulin (shown in red) and SMI 31 (shown in green) to confirm neuronal and axonal identity respectively.

**Video 6.** Long-term time-lapse of a stage 1 rat hippocampal neuron treated with 2µM CytoD, PFA fixed after 52 hrs., immunostained for βIII tubulin (shown in red) and SMI 31 (shown in green) to confirm neuronal and axonal identity respectively.

**Video 7.** Long-term time-lapse of a stage 1 rat hippocampal neuron treated with 5nM Taxol, PFA fixed after 52 hrs., immunostained for βIII tubulin (shown in red) and SMI 31 (shown in green) to confirm neuronal and axonal identity respectively.

**Video 8.** Long-term time-lapse of a stage 2 rat hippocampal neuron treated with DMSO, PFA fixed after 52 hrs., immunostained for βIII tubulin (shown in red) and SMI 31 (shown in green) to confirm neuronal and axonal identity respectively.

**Video 9.** Long-term time-lapse of a stage 2 rat hippocampal neuron treated with 2µM CytoD, PFA fixed after 52 hrs., immunostained for βIII tubulin (shown in red) and SMI 31 (shown in green) to confirm neuronal and axonal identity respectively.

**Video 10.** Long-term time-lapse of a stage 2 rat hippocampal neuron treated with 5nM Taxol, PFA fixed after 52 hrs., immunostained for βIII tubulin (shown in red) and SMI 31 (shown in green) to confirm neuronal and axonal identity respectively.

**Video 11.** Unprocessed (left) and background subtracted (right) 5 minute time-lapse (1 frame every 2 sec) of a EB3-tdTomato-transfected stage 2 control mouse cortical neuron,

**Video 12.** Unprocessed (left) and background subtracted (right) 5 minute time-lapse (1 frame every 2 sec) of a stage 2 mouse cortical neuron co-transfected with EB3-tdTomato and Cep120 shRNA.

**Video 13.** Unprocessed (left) and background subtracted (right) 5 minute time-lapse (1 frame every 2 sec) of a stage 2 mouse cortical neuron co-transfected with EB3-tdTomato and Cep120-GFP.

## Notes

### Competing Interest Statement

The authors have declared no competing interest.

